# Unsupervised physiological noise correction of fMRI data using phase and magnitude information (PREPAIR)

**DOI:** 10.1101/2022.02.18.480884

**Authors:** David Bancelin, Beata Bachrata, Saskia Bollmann, Pedro de Lima Cardoso, Pavol Szomolanyi, Siegfried Trattnig, Simon Daniel Robinson

## Abstract

Of the sources of noise which affect BOLD fMRI, respiration and cardiac fluctuations are responsible for the largest part of the variance, particularly at high and ultra-high field. Existing approaches to removing physiological noise either use external recordings, which can be unwieldy and unreliable, or attempt to identify physiological noise from the magnitude fMRI data. Data-driven approaches are limited by sensitivity, temporal aliasing and the need for user interaction. In the light of the sensitivity of the phase of the MR signal to local changes in the field stemming from physiological processes, we have developed an unsupervised physiological noise correction method which uses the information carried in both the phase and the magnitude of EPI data. Our technique, Physiological Regressor Estimation from Phase and mAgnItude, sub-tR (PREPAIR) derives time series signals which are sampled at the slice TR from both phase and magnitude images. It allows physiological noise to be captured without aliasing, and efficiently removes other sources of signal fluctuations which are not related to physiology, prior to regressor estimation. We demonstrate that the physiological signal time courses identified with PREPAIR not only agree well with those from external devices, but also retrieve challenging cardiac dynamics. The removal of physiological noise was as effective as that achieved with the most commonly used approach based on external recordings, RETROICOR. In comparison with widely used physiological noise correction tools which do not use external signals, PESTICA and FIX, PREPAIR removed more respiratory and cardiac noise and achieved a larger increase in tSNR at both 3 T and 7 T.

## 1. Introduction

Functional Magnetic Resonance Imaging (fMRI) using the Blood Oxygen Level Dependent (BOLD) (Ogawa et al., 1990) contrast has become the most powerful and ubiquitous method to investigate brain function during task processing and at rest. BOLD signals are corrupted, however, by non-white noise arising from scanner instability, head motion and physiological fluctuations related to the cardiac and respiratory cycles (Caballero-Gaudes and Reynolds, 2017; Liu, 2016; Lund et al., 2006; Reynaud et al., 2017). These decrease the temporal signal-to-noise-ratio (tSNR), limit the tSNR gain associated with increasing static magnetic field (Triantafyllou et al., 2005) and reduce sensitivity to BOLD activation (Biswal et al., 1996; Lund et al., 2006).

Respiration and cardiac-related fluctuations in EPI-based BOLD fMRI are responsible for most of the variance in grey matter and cerebrospinal fluid (CSF) (Bianciardi et al., 2009; Jezzard et al., 1993; Weisskoff et al., 1993; Windischberger et al., 2002). Susceptibility variations during breathing engender small changes in the magnetic field which lead to artifacts in the BOLD signal (Raj et al., 2001), as do signals related to the chest moving at the respiration frequency, which also translates into head motion (Birn et al., 2008; Krüger and Glover, 2001). Moreover, the pulsatile nature of the cardiac cycle causes variations in the blood flow as well as tissue movements which can contaminate the measured MR signal in voxels containing major blood vessels (Dagli et al., 1999), while the brainstem is particularly affected by physiological noise due to its proximity to the fourth ventricle and arteries (Harvey et al., 2008; Beissner et al., 2011; Matt et al., 2019).

Several methods have been developed to remove physiological perturbations and increase statistical power and statistical validity. Filtering approaches (Biswal et al., 1996; Chuang and Chen, 2001) and statistical models (Agrawal et al., 2020) perform well with fast fMRI but can fail when the frequency of the physiological noise exceeds the Nyquist frequency, which is generally 2/*TR_vol_*, where *TR_vol_* is the volume repetition time. Despite the more rapid acquisition allowed by simultaneous multi-slice (SMS) echo-planar imaging (EPI) (Barth et al., 2016; Feinberg et al., 2010; Feinberg and Setsompop, 2013; Setsompop et al., 2012), *TR_vol_* is still generally in excess of what would be needed to critically sample cardiac fluctuations, which is e.g. 400 ms for a heart rate of 75 bpm. However, slice time series can be reordered into temporal rather than spatial order to critically sample physiological noise (Frank et al., 2001). Alternative techniques involve the simultaneous acquisition of physiological signals during the fMRI experiment. RETROICOR (Glover et al., 2000; Harvey et al., 2008) models physiological noise using Fourier expansions of the physiological recordings while other approaches model the respiration and cardiac response functions of lower frequency fluctuations (Birn et al., 2008, 2006; Chang et al., 2009). However, external recordings increase the time overhead in subject handling and can be unreliable if the breathing belt becomes loose or slips (leading to signal loss) or is too tight (leading to a saturation in values) (Kasper et al., 2017), or, in the case of photoplephysmographs, if noise (e.g. high-frequency, power-line interference, baseline drift, motion artefact) corrupt the signal (Elgendi, 2012).

As physiological fluctuations are clearly identified in specific anatomical regions of the brain, many authors have explored the possibility of isolating cardiac and respiration signals directly from the fMRI data. Most common unsupervised data-driven methods generate physiological regressors using component analysis techniques (Behzadi et al., 2007; Beissner et al., 2014; Churchill and Strother, 2013; Perlbarg et al., 2007) of noise regions of interests (ventricles, brainstem, and major blood vessels or in the CSF and white matter) or the whole brain (Beall, 2010; Beall and Lowe, 2007; Salimi-Khorshidi et al., 2014; Thomas et al., 2002) to identify voxels dominated by physiological noise. For example, PESTICA (Beall, 2010; Beall and Lowe, 2007; Shin et al., 2016) correlates ICA components with a prior spatial physiological noise distribution to extract the component with highest correlation in each slice. By reordering the selected components into a single time series (Frank et al., 2001), PESTICA increases the sampling rate, enabling the recovery of aliased signals. Other methods like FIX (Salimi-Khorshidi et al., 2014) use prior information in existing trained datasets to automatically classify ICA components as noise, or can be trained through by-hand classification to improve performance. This process, and the need to check that the components identified by FIX as noise, makes the process onerous in studies with large numbers of subjects.

Most widely used physiological noise removal techniques only use magnitude fMRI data. However, data-driven physiological correction stands to benefit from the use of phase information, as the phase of the MR signal is highly sensitive to the changes in B_0_ which accompany respiration (Hagberg et al., 2012, 2008; Petridou et al., 2009). Phase images are not generally saved in fMRI experiments, but this information is available as the inherent counterpart of the magnitude in the complex MRI data and can be reconstructed and used for physiological noise correction. Indeed, there is a growing interest in the information carried by the phase in fMRI to increase the statistical power in fMRI analysis (Rowe, 2005), for dynamic distortion correction (Dymerska et al., 2018; Robinson et al., 2021), Quantitative Susceptibility Mapping (QSM) (Sun and Wilman, 2015) and functional QSM (Balla et al., 2014).

Using EPI phase poses some challenges, however: the need to find a solution to the phase-sensitive combination of RF coil signals, particularly at ultra-high field (Robinson et al., 2017), to unwrap phase images quickly and effectively and to remove unwanted signals from sources such as the cold-head helium pump which affect the phase signal to a larger extent than the magnitude (Hagberg et al., 2012). While a small number of methods have made limited use of phase – to identify Gaussian-distributed independent and identically distributed noise (Vizioli et al., 2021), for respiration (Cheng and Li, 2010; Le and Hu, 1996; Zahneisen et al., 2014), combined with ICA (Curtis and Menon, 2014) – none has taken full advantage of the sensitivity of phase to physiological fluctuations in a correction method for both respiratory and cardiac noise.

We propose a method which leverages phase and magnitude information, and obviates the need for external physiological signals and user interaction. This unsupervised, physiological data-free and automatic Physiological Regressor Estimation from Phase and mAgnItude sub-tR (PREPAIR) approach reveals both respiration and cardiac signals present in the fMRI time series for a wide range of TRs despite cardiac fluctuations being – considering the volume TR - critically sampled or undersampled. We adopt recently-developed approaches to coil combination (Eckstein et al., 2018) and unwrapping (Dymerska et al., 2021) combined with temporal reordering and slice filtering to derive physiological signals sampled at the slice repetition interval rather than the volume repetition interval. High-frequency fluctuations due to breathing motion and cardiac pulsatility are identified over nuisance signal variation. PREPAIR is compared with three commonly used physiological noise removal tools in both the cerebrum and the brainstem at 3 T and 7 T; RETROICOR using external physiological recordings (hereafter RETROICOR-EXT), and the ICA-based methods FIX and PESTICA at 3 T and 7 T.

## 2. Materials and Methods

This section comprises descriptions of subjects, MR measurements and external physiological recordings (Sections 2.1 and 2.2), the PREPAIR algorithm for estimating physiological noise regressors (Section 2.3), data processing steps (Section 2.4), and comparison with other methods (Section 2.5).

### 2.1. Study participants

All healthy volunteers participating to this study signed an informed consent form under a protocol approved by the Ethics Committee of the Medical University of Vienna. All were instructed to rest with eyes closed and to remain as still as possible. Ten subjects took part in the 3 T study (5 females, age range 25 – 43 years old, mean age 31.1 ± 6.6 years old) and five subjects in the 7 T study (one female, four males, age range 26 – 38 years old, mean age 31.3 ± 4.6 years old).

### 2.2. Measurements

PREPAIR was tested on 3 T data with whole brain coverage and, to explore the dependence of the method on field strength and to analyze brain structures particularly prone to physiological noise, on 7 T data with a sagittal acquisition covering the brainstem. Protocols with a range of TRs and multi-band factors were used in each study.

fMRI data were acquired using a multi-band accelerated 2D-EPI pulse sequence (Xu et al., 2013) developed at the Centre for Magnetic Resonance Research, University of Minnesota, Minneapolis U.S.A. (CMRR). The experiment comprised runs with four protocols with a range of TR, TE, flip angle (FA), multi-band factor (MB), number of slices (N), number of repetitions (NR), slice thickness (ST) and GRAPPA acceleration factor detailed below and in Table 1.

**Table 1:**
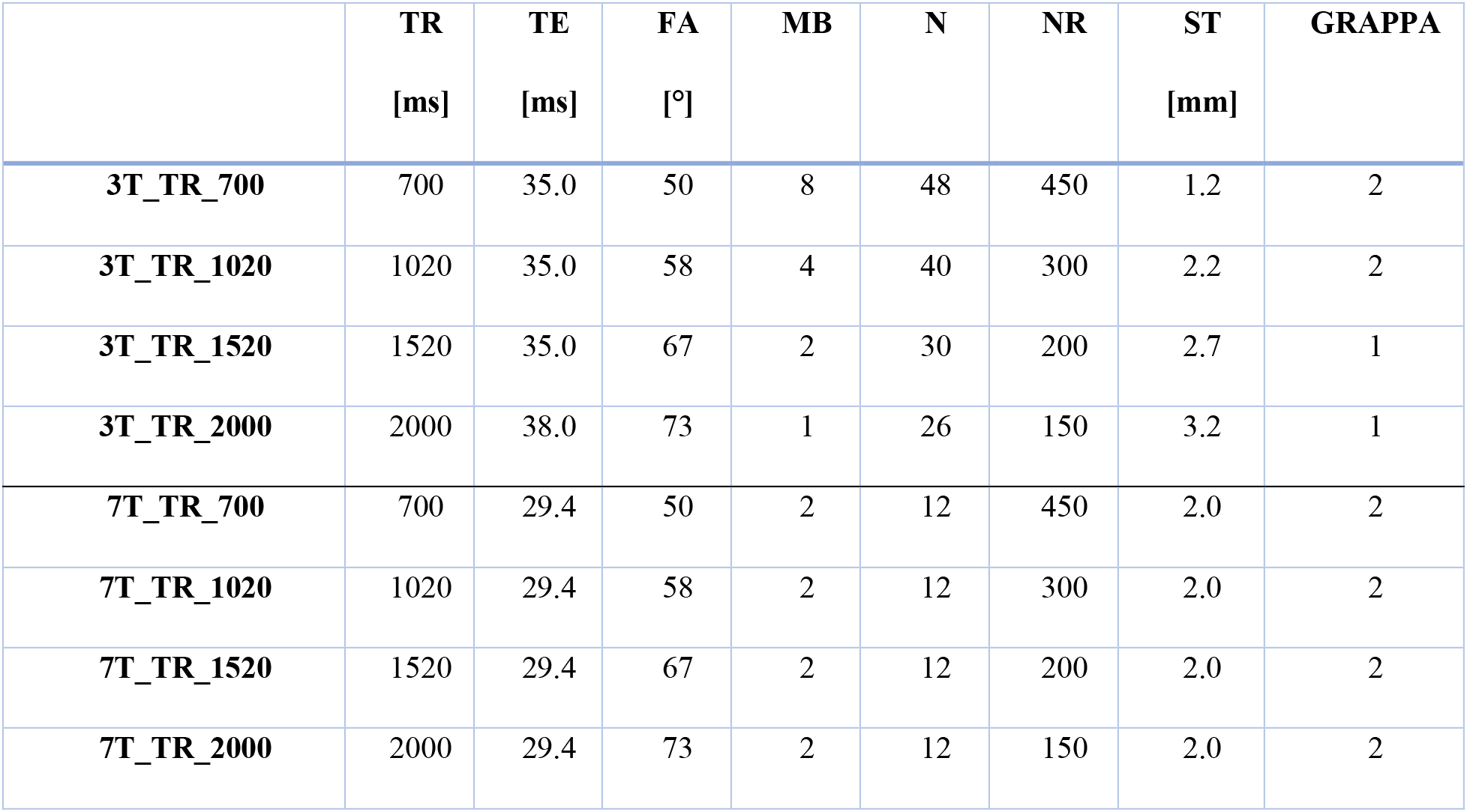
Acquisition parameters for the 3 T and 7 T studies. FA = flip angle, MB = multiband acceleration factor, NR = number of repetitions and ST = slice thickness. First column indicates protocol names.

Each fMRI measurement was preceded by a low resolution, monopolar dual-echo gradient-echo prescan which was used to calculate the phase offset for each RF coil using ASPIRE (Eckstein et al., 2018). To generate channel-combined EPI phase images, these phase offsets were subtracted from the phase of each channel of the EPI data for each time point, online in the image reconstruction environment, prior to calculating the angle of the magnitude-weighted complex sum over channels (Bachrata et al., 2018).

#### 3 T study

Whole brain axial MRI data were acquired with a 3 T Siemens PRISMA (Siemens, Erlangen, Germany) scanner with a 64-channel head coil. Neck coils were turned off. Slices were acquired parallel to the AC-PC plane with a matrix size of 128*128, FOV=210 mm * 210 mm (1.8 mm*1.8 mm in-plane resolution), distance factor of 20%, phase encoding direction: posterior-anterior. Protocols parameters are summarized in Table 1.

#### 7 T study

Sagittal slices were acquired with a Siemens MAGNETON 7T+ scanner with a 32-channel head coil with a matrix size of 128*128, FOV=220 mm * 220 mm (1.7 mm*1.7 mm* 2.0 mm spatial resolution), distance factor of 30%, phase encoding direction: posterior-anterior (see Table 1 for other protocol parameters and naming).

For comparison, physiological measurements of respiration and cardiac were made using arespiratory belt and an optical plethysmograph respectively. Physiological Monitoring Unit (PMU) signals were recorded by the CMRR sequence at 1/400 Hz intervals, along with scanner trigger signals.

### 2.3. Estimation of the physiological regressors

The central aim of the PREPAIR approach is to generate two scalar signals sampled at *TR_slice_* (rather than *TR_vol_*) – one from the magnitude and one from the phase of the EPI data – and select from the two that which better describes respiration and cardiac fluctuations. These are used to generate physiological noise regressors for correction of the magnitude EPI data. The process consists of the following steps (illustrated in Figure 1, in which the letters below appear):

a. Voxels with high signal-to-noise ratio (SNR) were identified using the criteria that their mean intensity over time in the magnitude was above 20% of the robust maximum defined as the 98^th^ percentile of the values in the magnitude data over all voxels (Aslan et al., 2019).
b. High SNR voxel values were averaged over each slice. For the phase, a weighted mean was used, in which weights were the corresponding magnitude values. This yielded a magnitude and phase time series for each slice, sampled at *TR_vol_*.
c. The dependence of the magnitude and phase values on imaging slice, which stems from different coverage and tissue composition, was eliminated by fitting and subtracting a 3^rd^ order polynomial to each slice time series.
d. For phase and magnitude separately, signals derived from each slice were combined according to the acquisition order of each contributing slice. For SMS data, the signal in simultaneously acquired slices was averaged and the number of slices was reduced to NS = N/MB. This resulted in a magnitude and phase time series with length NR*NS and sampling rate NS/TR.
e. A ‘prefiltering’ step, consisting of a bandpass filter with typical ranges of [9 – 30] cpm (cycle per minute) for respiration and [42 – 96] bpm for cardiac, was applied to select physiological noise contributions to the magnitude and phase signals.
f. The fundamental frequencies of the respiratory and cardiac noise, f_R_ and f_C_ respectively, were then identified with an iterative procedure in which the frequency with the highest power in the respective power spectrum was compared with signals with frequencies unrelated to physiological noise. In order not to remove signal of interests, unrelated physiological noise frequencies were removed by setting their power to zero rather than applying a bandpass filter to the time series.
g. The respiratory and cardiac frequency spectra were then reduced in width by applying a bandpass filter to the time series to retain only frequencies within a certain range around the fundamental frequencies.
h. To determine whether the PREPAIR-magnitude or PREPAIR-phase physiological time series better represented physiological fluctuations in the data, the variance improvement after separately removing the respiratory and cardiac noise by linear regression was estimated and compared for both magnitude-derived and phase-derived signals. This led to the selection of either the PREPAIR-magnitude or PREPAIR-phase time series for respiratory-related fluctuations, and the same for cardiac-related fluctuations.
i. Slice-wise physiological noise regressors were modeled from the PREPAIR respiratory and cardiac time series as sine and cosine functions of the two first harmonics of the cardiac and respiratory cycle using the AFNI version of RETROICOR-EXT.

**Figure 1:**
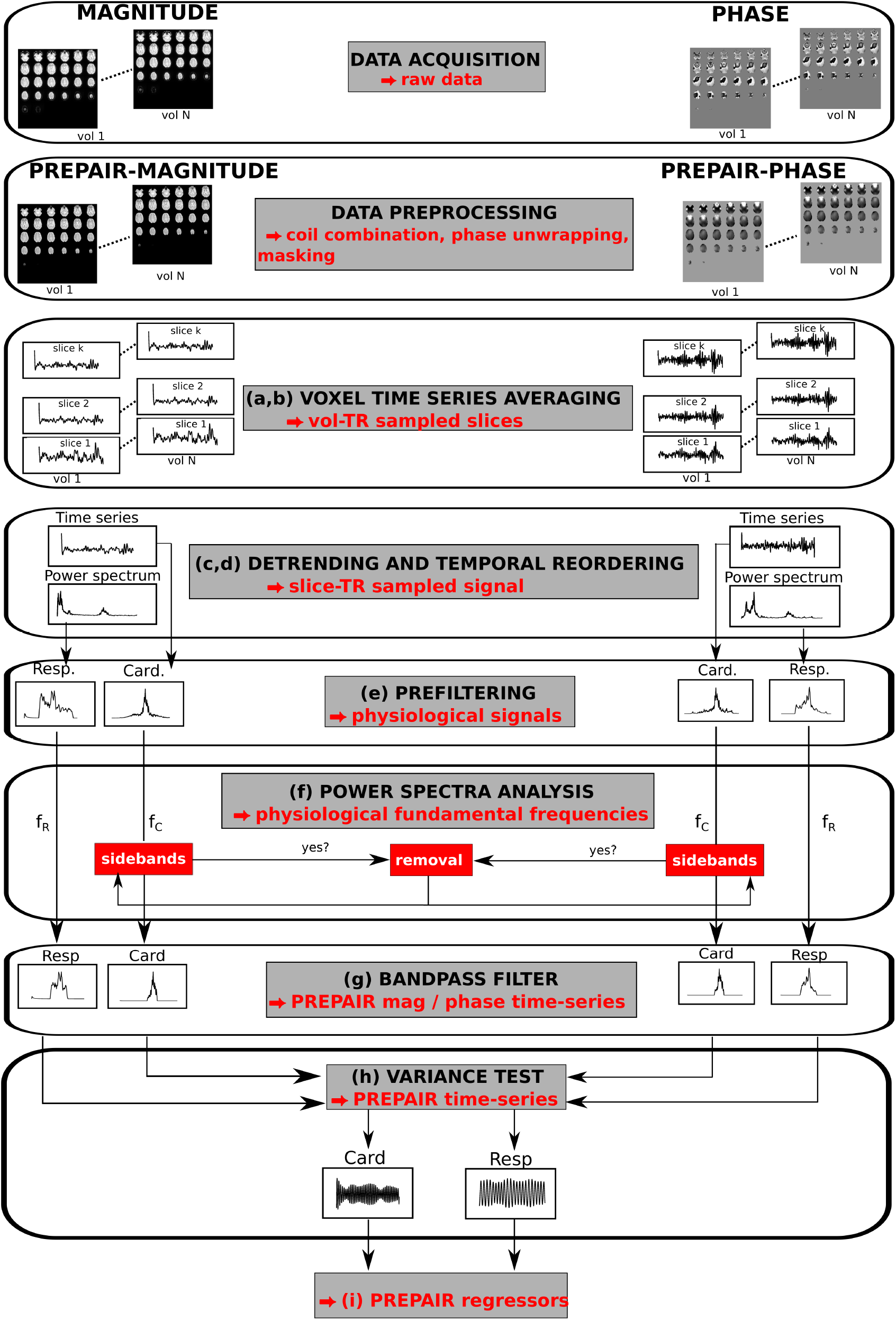
Steps in generating PREPAIR regressors (letters in brackets refer to steps described in Section 2.3). Raw magnitude and phase were preprocessed (for magnitude, with masking, for phase, with coil combination, unwrapping and masking). Magnitude and phase slices were averaged (a,b) then detrended and temporally reordered to yield slice TR sampled signals (c,d) from which respirators and cardiac time series were prefiltered (e). After deriving the corresponding physiological fundamental frequencies (f), the narrowed power spectra (g) underwent a variance test to choose between magnitude and phase physiological signals (h) to create the PREPAIR regressors (i).

### 2.4. Data processing

Phase wraps in raw phase images were removed by spatially unwrapping using ROMEO (Dymerska et al., 2021).

In the 3 T study, an image mask was generated using AFNI (Cox, 1996; Cox and Hyde, 1997), and applied to constrain analysis to the brain. In the 7 T study, no mask was used in the physiological noise identification or in the magnitude image correction, but analysis was constrained to a brainstem mask which included voxels in the medulla neighboring the spinal cord (which would be discarded with AFNI) by selecting voxels exceeding an intensity of 3000.

Data processing and analysis were performed using Matlab 2018b (The Mathworks Inc., Natick, MA, USA).

### 2.5. Comparison with other methods

Among the existing physiological noise correction methods, we chose to compare PREPAIR with RETROICOR using external physiological recordings (RETROICOR-EXT), PESTICA, which is ICA-based but shares the similarity with PREPAIR that it also generates a slice TR signal, and FIX, another widely used ICA-based tool.

For RETROICOR-EXT, physiological signals recorded with the CMRR sequence were processed with the PhysIO Toolbox to model physiological noise regressors using RETROICOR with respect to the middle slice (Kasper et al., 2017).

PESTICA (Shin et al., 2016) (version 5.2) computes physiological estimators which can be filtered either via an interactive window inviting the user to zoom in to an appropriate region (supervised mode) or by using a default window of [48 – 85] bpm for cardiac and [10 – 24] cpm for respiration (unsupervised mode). Because of the large number of protocols and participants in our study, our analysis was performed in unsupervised mode, in which the frequencies of the respiratory and cardiac estimators ranged between [10 –24] cpm and [45 – 85] bpm, respectively.

For FIX (version 1.06.15), MELODIC of FMRIB’s Software Library (FSL version 4.15; www.fmrib.ox.ac.uk/fsl) was performed, with the number of components into which to separate the data set to 30. These were classified by FIX using the HCP7T_hp2000.Rdata training file. This was chosen because the spatial resolution was most similar to our spatial resolution. Due to the large number of tests performed in our study, we decided to run FIX in unsupervised mode, in which the threshold of good versus bad components was set to be the same for all subjects. This value was set to 50 after checking for a small number of subjects that resting state networks were not included in the list of components to be removed. This led to the identification of ~ 6 components/subject to for the 3 T study and ~2 components/subject for the 7 T study to be filtered out.

#### 2.5.1. Correlation with the physiological recordings

Because the PREPAIR physiological time series were bandpass-filtered such that the window of their power spectra was centered on their corresponding fundamental frequency with a range of frequencies around it (as described in Section 2.3, step g), we applied the same bandpass filter to the respiratory and cardiac waveforms derived from the different methods to narrow their power spectrum to the same window size as PREPAIR’s, to ensure a fair comparison. All correlations in Section 3.1 accounted for lags between the physiological noise from the fMRI data and the physiological recordings by measuring the cross correlation between those time series. Although FIX provides the list of components removed from the fMRI data, no correlations were computed, as that would have required a manual inspection of the classified components to only include those which were physiological noise-related; a process which is both time consuming and subjective.

#### 2.5.2. Spectral Analysis

To facilitate the visualization of the physiological noise present in 4D time series of the corrected magnitude images in Section 3.2, data were converted into a single time series sampled at the slice TR by first averaging over the voxels in each slice, then reordering those time series according to the slice acquisition times. We then performed a spectral analysis of these time series using the Chronux toolbox (Bokil et al., 2010) to produce spectra of frequencies of signals varying in average sliding windows of 45s and a step-size of 4s (Supplementary Information sFig. 2 (the first five subjects of the 3 T study), sFig. 3 (the last five subjects of the 3 T study) and sFig. 4 (7 T study)).

To assess the image correction efficiency of each method presented in Table 3 and Table 4, we calculated the proportion of noise removed in the respiratory and cardiac bands (including the second harmonic when present) of the uncorrected magnitude data by computing the relative change in power for each voxel between the uncorrected and corrected magnitude, and then by averaging over each slice. When negative (i.e. the correction added noise), this proportion was set to zero and added to the data. Zero values of the proportion mean that the correction was either non-effective or added noise, and values of one indicate that the level of physiological noise decreased by 100%. To determine which method preserved the integrity of the signal outside the respiration and cardiac areas, we also included the proportion of power fluctuation, i.e. the change in the power signal (increase or decrease) after correction. Contrary to the previous step, negative values of the proportion were not set to zero. Values close to zero would indicate that a method was highly effective in preserving the original signal.

#### 2.5.3. tSNR improvement in anatomical regions

To assess tSNR improvement in regions of the brain typically affected by physiological fluctuations, we performed a statistical analysis in manually-drawn 3D ROIs in the visual and insular cortices in the 3 T study (subjects 1, 2, 4, 7 and 10), and the brainstem in the 7 T study.

### 2.6. Data and code availability statement

All source data are publicly available at the Harvard Dataverse (https://dataverse.harvard.edu/dataverse/prepair). The package PREPAIR is available at (https://github.com/daveB1978/PREPAIR).

## 3. Results

In this section, we illustrate the physiological signals generated with PREPAIR and compare physiological noise correction between PREPAIR, RETROICOR-EXT, PESTICA and FIX. In Section 3.1, the quality of physiological waveforms obtained with each method is compared with that from the external signals. The efficacy of the noise removal with each method is shown in Section 3.2 and its effectiveness in the improvement of the magnitude image in Section 3.3.

Because all intermediate data needed for this assessment are not available with FIX (e.g. slice TR-sampled physiological regressors), the comparison between PREPAIR and FIX was limited to the analysis of the corrected magnitude image (Section 3.2).

### 3.1. Physiological noise identification

#### Illustration of the PREPAIR algorithm

The effect of the steps detailed in Section 2.3 for physiological noise identification is illustrated in Figure 2 with the phase data of one subject (S6, 3T_TR_700). Panel A shows the slice TR sampled phase signal (top). Before removing slice effects, both slow fluctuations (blue square) and faster fluctuations (slice groups, green arrows) are apparent. The Fourier spectrum (bottom) is dominated by three peaks: f_pump_ at 0.13 Hz, a disturbance related to the cold-head helium pump affecting the flow of the gradient cooling water (as described in Hagberg et al. (2012)), f_R_, the respiratory fundamental at 0.25 Hz and f_1/TR_ at ~1.43 Hz caused by slice reordering. Slice effects were removed by detrending (Section 2.3, c and d) (panel B), reducing the power of f_1/TR_ and revealing rapid variations of cardiac origin (red arrows). In panel C, respiratory fluctuations (left) were bandpass filtered from the power spectrum (Section 2.3, e), resulting in a correlation of 0.78 with the external signal for the case illustrated. The cardiac fluctuations (right) extracted from the time series by applying a 2^nd^ order Butterworth bandpass filter had a correlation value of 0.37 with the external signal. This Butterworth filter includes frequencies beyond the chosen interval so that unusual cardiac rhythms can be identified. In this cardiac power spectrum, we observe multiple peaks: the cardiac fundamental f_C_ and sidebands caused by f_pump_ located at S_pump_ = f_1/TR_ ± f_pump_ and by f_R_ located at S_R_ = f_1/TR_ ± f_R_, respectively (the upper sideband does not appear as it lies outside the cardiac range). Here, f_C_ was well below S_pump_ and S_R_ which would be wrongly identified as the fundamental. After filtering out all sidebands (Section 2.3, f), f_C_ was correctly identified (panel D) which improved the correlation with the external signals to 0.49. In panel E, the intervals of the respiratory and cardiac frequencies were reduced to ranges around f_R_ and f_C_ (Section 2.3, g). These ranges were established by computing the average standard deviation of the respiration and cardiac rates in the signals obtained from the external signals over all subjects, protocols and field strengths, which yielded 2.5 cpm and 6.5 bpm for respiration and cardiac, respectively. Frequencies outside the ranges f_R_ ± 2.5 cpm and f_C_ ± 6.5 bpm were eliminated with a bandpass filter, leading to correlation for respiration and cardiac with the external signals of 0.97 and 0.81 respectively.

**Figure 2:**
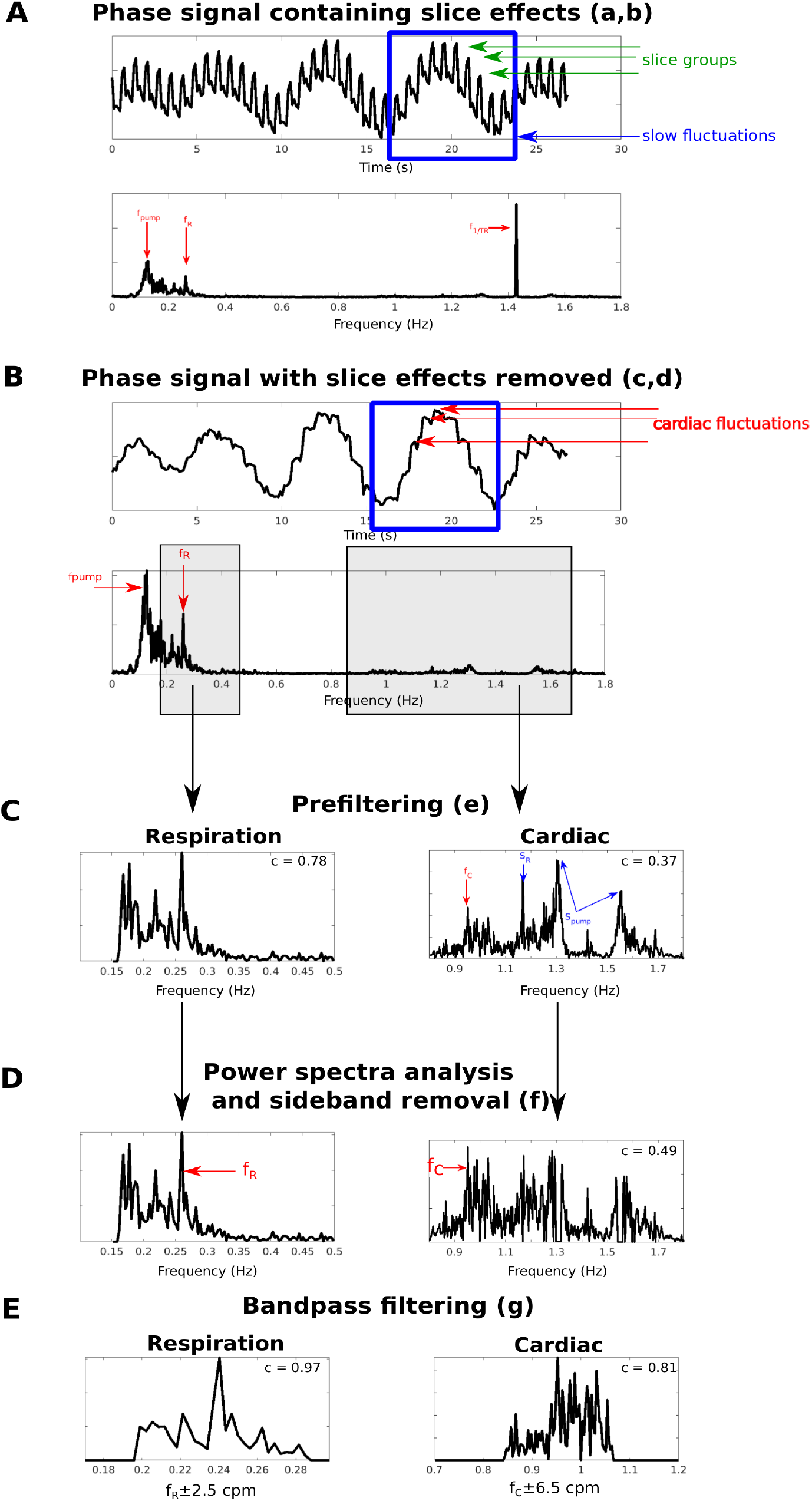
Illustration of physiological noise identification in the slice TR sampled phase signal of subject S6, 3T_TR_700 (A), which is dominated by periodic slice effects (green arrows) sampled at f_1/TR_, and slow fluctuations (blue square) comprising respiration (sampled at 1/f_R_) and cold head pump (sampled at 1/f_pump_). After removing the slice effects (B), physiological fluctuations were prefiltered from the phase signal (C) and unrelated physiological frequencies (sidebands S_R_ and S_pump_) were removed to find the correct cardiac fundamental frequency f_C_ (D). Respiratory and cardiac signals were bandpass filtered around f_R_ and f_C_ respectively. Correlations improvement with the external signals are indicated for each step of the algorithm.

#### Comparison of the physiological waveforms

Table 2 lists the mean correlation over subjects of PREPAIR-magnitude and PREPAIR-phase waveforms with the external signals and the proportions of cases in which the PREPAIR-magnitude and PREPAIR-phase regressors were selected for respiratory and cardiac noise. For the 3 T study, PREPAIR-phase matched extremely well the external signals and modeled respiration fluctuations better than PREPAIR-magnitude for all subjects and protocols. For cardiac, correlation values with PREPAIR-phase were lower, and were identified as the best regressor in 40% of cases for 3T_TR_700 and 3T_TR_1020 and 60% for 3T_TR _1520 and 3T_TR_2000 protocols. For the 7 T study, PREPAIR-phase also provided a better description of respiration-related fluctuations for all subjects and protocols. For cardiac, the PREPAIR-magnitude waveform was used to build the cardiac regressors in all subjects and protocols.

**Table 2:** PREPAIR-phase and -magnitude correlation with the external signals, and proportion of PREPAIR-phase and - magnitude (phase vs magnitude) selected by the algorithm for deriving the PREPAIR regressors. PREPAIR-phase modeled respiratory fluctuations better than PREPAIR-magnitude for both 3 T and 7 T studies and all PREPAIR-phase signals were used for deriving respiratory regressors. For cardiac, PREPAIR-phase and -magnitude contributed equally for the 3 T study over all protocols, whereas for the 7 T study, PREPAIR-magnitude was chosen in almost all protocols (except for 7T_TR_1020).

| 3T |  | 3T_TR_700 | 3T_TR_1020 | 3T_TR_1520 | 3T_TR_2000 |
| --- | --- | --- | --- | --- | --- |
| **Respiration** | Correlation:<br>phase | $0.97 \pm 0.02$ | $0.97 \pm 0.03$ | $0.96 \pm 0.04$ | $0.95 \pm 0.03$ |
| | Correlation:<br>magnitude | $0.66 \pm 0.21$ | $0.60 \pm 0.19$ | $0.63 \pm 0.25$ | $0.18 \pm 0.28$ |
|  | Phase vs<br>magnitude | 100%:0% | 100%:0% | 100%:0% | 100%:0% |
| **Cardiac** | Correlation:<br>phase | $0.72 \pm 0.28$ | $0.61 \pm 0.36$ | $0.73 \pm 0.25$ | $0.43 \pm 0.28$ |
| | Correlation:<br>magnitude | $0.90 \pm 0.10$ | $0.75 \pm 0.23$ | $0.68 \pm 0.24$ | $0.25 \pm 0.24$ |
|  | Phase vs<br>magnitude | 40%:60% | 40%:60% | 60%:40% | 60%:40% |
| 7T |  | 7T_TR_700 | 7T_TR_1020 | 7T_TR_1520 | 7T_TR_2000 |
| **Respiration** | Correlation:<br>phase | $0.76 \pm 0.24$ | $0.93 \pm 0.02$ | $0.93 \pm 0.04$ | $0.93 \pm 0.02$ |
| | Correlation:<br>magnitude | $0.75 \pm 0.34$ | $0.78 \pm 0.22$ | $0.59 \pm 0.31$ | $0.33 \pm 0.44$ |
|  | Phase vs<br>magnitude | 100%:0% | 100%:0% | 100%:0% | 100%:0% |
| **Cardiac** | Correlation:<br>phase | $0.79 \pm 0.11$ | $0.67 \pm 0.33$ | $0.70 \pm 0.18$ | $0.59 \pm 0.32$ |
| | Correlation:<br>magnitude | $0.89 \pm 0.03$ | $0.85 \pm 0.06$ | $0.80 \pm 0.09$ | $0.80 \pm 0.11$ |
|  | Phase vs<br>magnitude | 0:100% | 20%:80% | 0%:100% | 0%:100% |

Given the important contribution of the phase and magnitude data in the process of noise identification, we compared, in Figure 3, the correlation of the PREPAIR respiratory and cardiac waveforms selected by the algorithm with that of PESTICA (all PESTICA estimators are presented in Supplementary Information sFig. 8 and sFig. 9). For the 3 T and 7 T studies, PREPAIR performed significantly better (*: p<0.05; **: p<0.01; ***: p<0.001) than PESTICA except for 7T_TR_700 for respiration.

**Figure 3:**
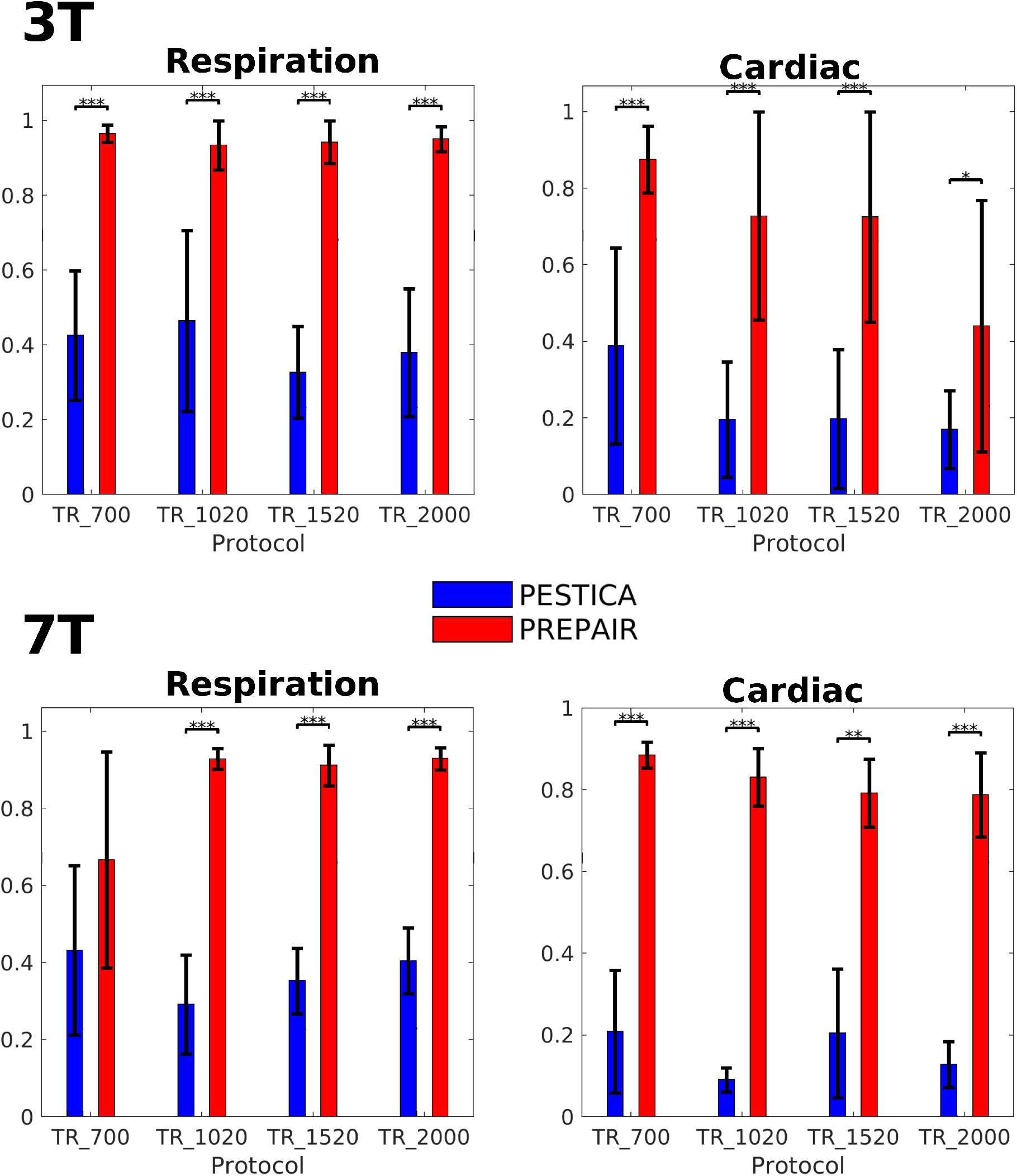
Correlation of PESTICA (blue) and PREPAIR (red) with the externals signals across subjects for respiration and cardiac for the 3 T study (top) and 7 T study (bottom). Most mean correlations of PREPAIR are significantly (*: p<0.05; **:p<0.01; ***: p<0.001) above those of PESTICA.

The results of this section demonstrate that PREPAIR can accurately detect respiratory and cardiac fluctuations in the fMRI data. The respiratory and cardiac waveforms selected by the algorithm were used to produced physiological noise regressors for image correction.

### 3.2. Physiological noise removal

#### Comparison of uncorrected versus corrected power spectra

The effectiveness of noise removal with the methods under consideration was investigated by performing a spectral analysis of the magnitude data prior to and following correction. In the analysis below, we compare the power reduction of physiological noise bands with the three methods for three selected subjects in the 3 T study who had low (top: S10, 3T_TR_700), normal (middle: S5, 3T_TR _1520) and high (bottom: S4, 3T_TR_1020) cardiac heart rate. In Figure 4, the variation of the respiratory and cardiac fluctuations over time obtained with the external recordings is indicated with a black and red arrows (two if the 2^nd^ harmonic is visible) in the uncorrected magnitude image (left panel) respectively. Spectrograms were scaled to the same minimum and maximum power expressed in decibels (color scale).

a. In the top panel, showing results for a subject with a slow pulse, the respiratory noise was close to 0.3 Hz and the cardiac 1^st^ and 2^nd^ harmonics were at ~0.6 Hz and 1.2 Hz. The respiratory removal was as effective with PREPAIR and RETROICOR-EXT, while in PESTICA the correction was partial, and in FIX non-effective. The contributions of the 1^st^ and 2^nd^ cardiac harmonics were completely removed by correction with PREPAIR and RETROICOR-EXT while PESTICA and FIX only removed the 2^nd^ harmonic.
b. In the middle panel, showing results for a subject with a typical pulse, the respiratory noise lies near 0.3 Hz and the cardiac 1^st^ harmonic around 1.0 Hz. Concerning the respiration, PREPAIR slightly outperformed RETROICOR-EXT, for which some islets of power remain in the spectrograms (black arrows with label 1), and PESTICA and FIX added noise. As for the cardiac noise, remaining power is visible in the spectrogram of PREPAIR compared to RETROICOR (red arrows with label 2), but neither FIX nor PESTICA filtered this noise out.
c. In the bottom panel, showing results for a subject with a fast pulse, the respiratory noise peak was around 0.3 Hz and the cardiac 1^st^ harmonic around 1.6 Hz. Physiological noise removal was only successful with RETROICOR-EXT and PREPAIR, albeit with one islet of residual power (red arrow with label 3) in PREPAIR’s spectrogram.

**Figure 4:**
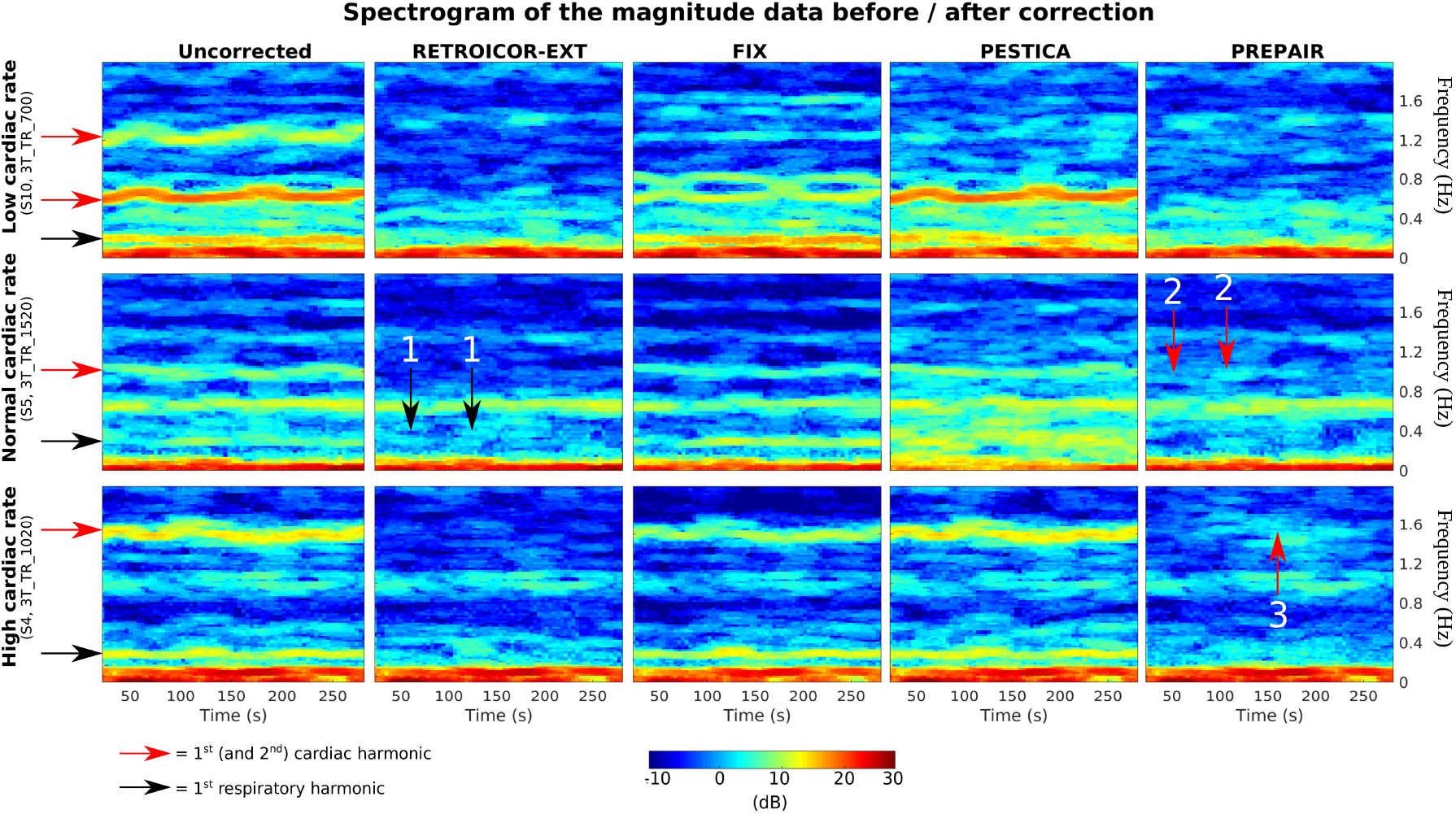
Spectrograms of uncorrected (left) versus corrected magnitude data. Frequencies (y-axis) were truncated at 1.7 Hz and rows were scaled to the same power expressed in decibels (color scale). Horizontal black and red arrows indicate the 1^st^ (and 2^nd^ when applicable) harmonic of the respiratory and cardiac noise, respectively, obtained with the external signals. PREPAIR showed effective noise removal in comparison with the other methods, even with unusual cardiac rates (top and bottom). Numbered vertical arrows show islets of remaining power after noise correction.

To estimate the overall performance of each method after physiological noise correction, we computed the mean proportion of physiological noise removed and of fluctuation outside the physiological noise regions. Figure 5 shows the mean proportion of physiological noise removed in the 3 T (top panel) and 7 T (bottom panel) studies across subjects. For the 3 T study, PREPAIR gave a correction which was as effective as RETROICOR-EXT, slightly less effective with FIX for some protocols, but more effective than PESTICA. For the 7 T study, PREPAIR performed better than PESTICA and FIX for all protocols, and similarly to RETROICOR-EXT. Table 3 compiles the mean and standard deviation (in percentage) of the total noise removed by each method over all subjects and protocols for the 3 T study. For respiration (first row), PREPAIR (32.7%) was as effective as RETROICOR-EXT (33.3%) and slightly less than FIX (35.7%), with the performance of PESTICA (16.8%) being significantly lower (p<0.001). For cardiac (second row), PREPAIR (25.9%) performed similarly to RETROICOR-EXT (25.2%), significantly better than PESTICA (15.7%: p<0.001). However, FIX (35.2) removed significantly more cardiac noise than PREPAIR (p<0.001). Table 4 shows similar results for the 7 T study. For respiration, PREPAIR (31.7%) was as effective as RETROICOR-EXT (30.6%), and removed significantly more respiratory signals than FIX (11.3%: p<0.001) and PESTICA (13.7%: p<0.001). For cardiac, PREPAIR (27.2%) also performed significantly better than FIX (12.8%: p<0.001) and PESTICA (12.8%: p<0.001), and similarly to RETROICOR-EXT (27.7%).

**Figure 5:**
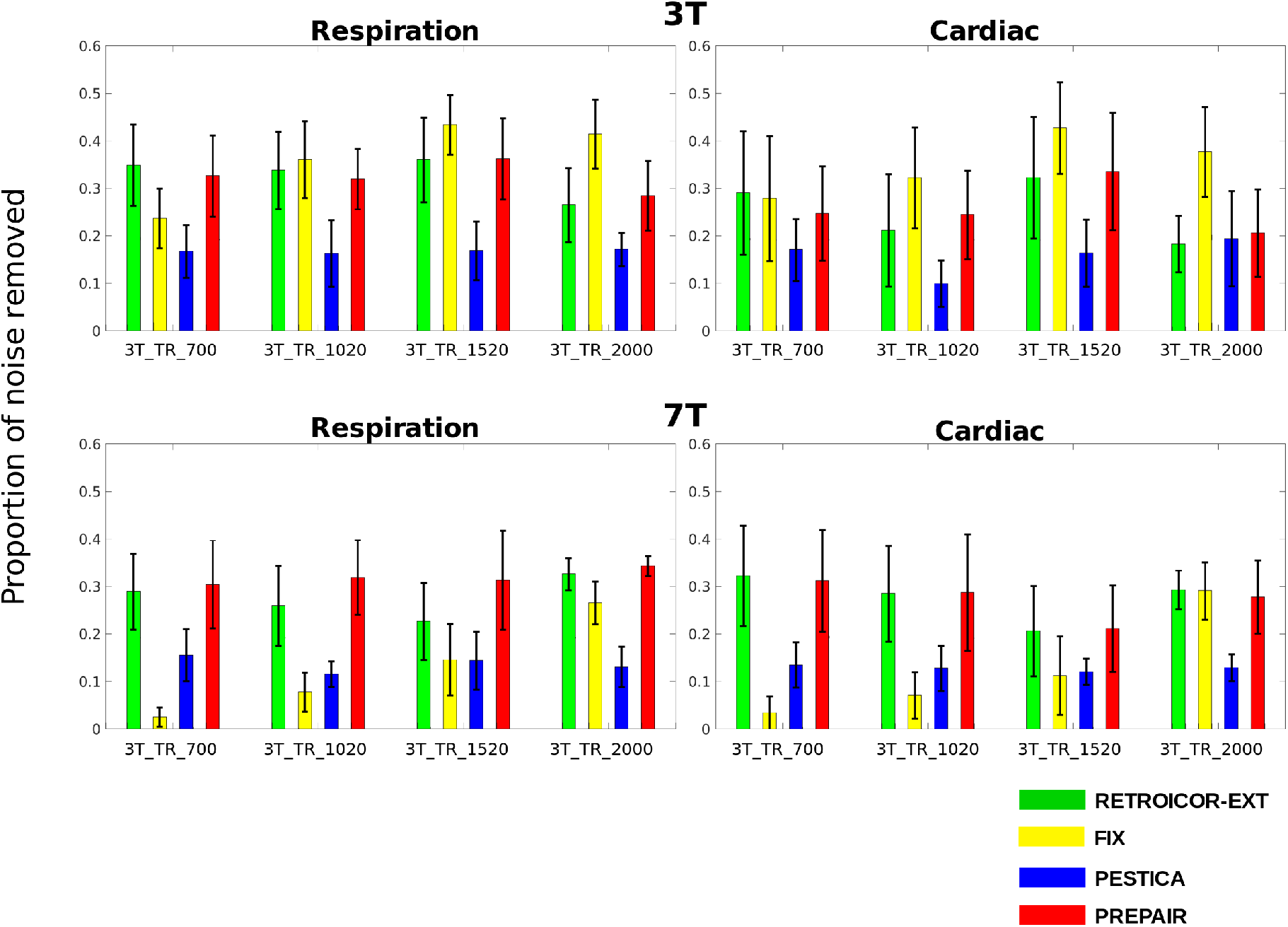
Proportion of respiratory and cardiac noise removed by each method and for each protocol across subjects. Physiological noise correction performed with PREPAIR was as effective as that of RETROICOR-EXT but more effective than FIX at 7 T only and PESTICA at both 3 T and 7 T.

**Table 3:** Mean and standard deviation of the percentage of physiological noise removed (two first rows) in the physiological bands and power fluctuation (last row) outside these bands, by each method over all subjects and protocols for the 3 T study. PREPAIR was slightly less effective than FIX in decreasing power in the respiratory band but significantly more effective than PESTICA (***: p<0.001 and similar to RETROICOR-EXT. Cardiac noise reduction with PREPAIR was significantly less than FIX (p<0.001). The power in other spectral regions (Power fluctuation) with PREPAIR was changed significantly less than the FIX (p<0.001) and PESTICA (p<0.01).

| 3 T | RETROICOR-EXT | FIX | PESTICA | PREPAIR |
| --- | --- | --- | --- | --- |
| **% of noise removed:**<br>**Respiration** | 33.3 ± 8.8 | 35.7 ± 10.4 | 16.8 ± 5.6 *** | 32.7 ± 7.9 |
| **% of noise removed:**<br>**Cardiac** | 25.2 ± 12.3 | 35.2 ± 11.9*** | 15.7 ± 7.9 *** | 25.9 ± 11.0 |
| **Power fluctuation**<br>**(%)** | 1.4 ± 2.7 | 28.4 ± 11.8*** | 3.3 ± 5.7** | 1.9 ± 5.0 |

**Table 4:** Mean and standard deviation of the percentage of physiological noise removed (two first rows) in the physiological bands and power fluctuation (last row) outside these bands, by each method over all subjects and protocols for the 7 T study. For respiration, PREPAIR was significantly more effective in decreasing power in the respiration band than FIX and PESTICA (p<0.001), and similar to RETROICOR-EXT. Cardiac noise with PREPAIR was comparable to that of RETROICOR-EXT, and significantly more effective than FIX and PESTICA (p<0.001). The change in power in other spectral regions (Power fluctuation) with PREPAIR was slightly higher to that of RETROICOR-EXT and PESTICA but significantly more effective than FIX (p<0.001).

| 7 T | RETROICOR-EXT | FIX | PESTICA | PREPAIR |
| --- | --- | --- | --- | --- |
| % of noise removed:<br>Respiration | 30.6 ± 7.8 | 11.3 ± 9.6*** | 13.7 ± 4.7*** | 31.7 ± 7.9 |
| % of noise removed:<br>Cardiac | 27.7 ± 9.3 | 12.8 ± 11.5*** | 12.8 ± 3.6*** | 27.2 ± 10.0 |
| Power fluctuation<br>(%) | 1.1 ± 3.3 | -27.8 ± 23.9*** | 1.5 ± 5.4 | 2.5 ± 2.6 |

A reliable physiological noise correction method should be effective in removing the narrowband physiological noise and leave signal outside these regions unchanged. To assess the extent to which all methods modified signal power outside the physiological bands, we estimated the proportion of power fluctuation outside physiological noise regions. For the 3 T study (last row of Table 3), PREPAIR only altered the magnitude signal by 1.9%, comparable to RETROICOR-EXT (1.4%), which was significantly less than FIX (28.4%, p<0.001) and PESTICA (3.3%, p<0.01). For the 7 T study (last row of Table 4), PREPAIR (2.5%), RETROICOR-EXT (1.1%) and PESTICA (1.5%) all reduced the power fluctuation to similar level, whereas the change of power outside the physiological bands was significantly larger with FIX (−27.8%: p<0.001).

This section demonstrates that PREPAIR effectively removed physiological fluctuations from magnitude time series data while preserving signal of interests, which is crucial for the detection of task-driven voxels.

### 3.3. tSNR improvement in anatomical regions

tSNR increases in the insular cortex, visual cortex and brainstem for S2, TR_700 are illustrated in the left panel of Figure 6. The improvement in tSNR was similar with RETROICOR-EXT and PREPAIR, and larger than PESTICA, particularly in the brainstem. In the assessment over all subjects, shown in the boxplots in the right part of the same figure, it is apparent that the mean tSNR increase with PESTICA was less than 5% for all protocols, with significant differences between PESTICA and PREPAIR in all protocols and anatomical regions. The mean for RETROICOR-EXT and PREPAIR was above 5% in all cases. In the 3 T study, RETROICOR-EXT was slightly more effective than PREPAIR in the insular cortex (by a few percent) for all protocols. In the visual cortex, tSNR gain with PREPAIR was slightly lower for 3T_TR_700 and 3T_TR_1020, comparable to RETROICOR-EXT for 3T_TR_1520 and slightly better than RETROICOR-EXT for 3T_TR _2000. In the brainstem, 50^th^ percentiles of PREPAIR were always similar to those of RETROICOR-EXT for all protocols, although 75^th^ percentiles of RETROICOR-EXT for 7T_TR_700, 7T_TR_1520 and 7T_2000 were slightly higher.

**Figure 6:**
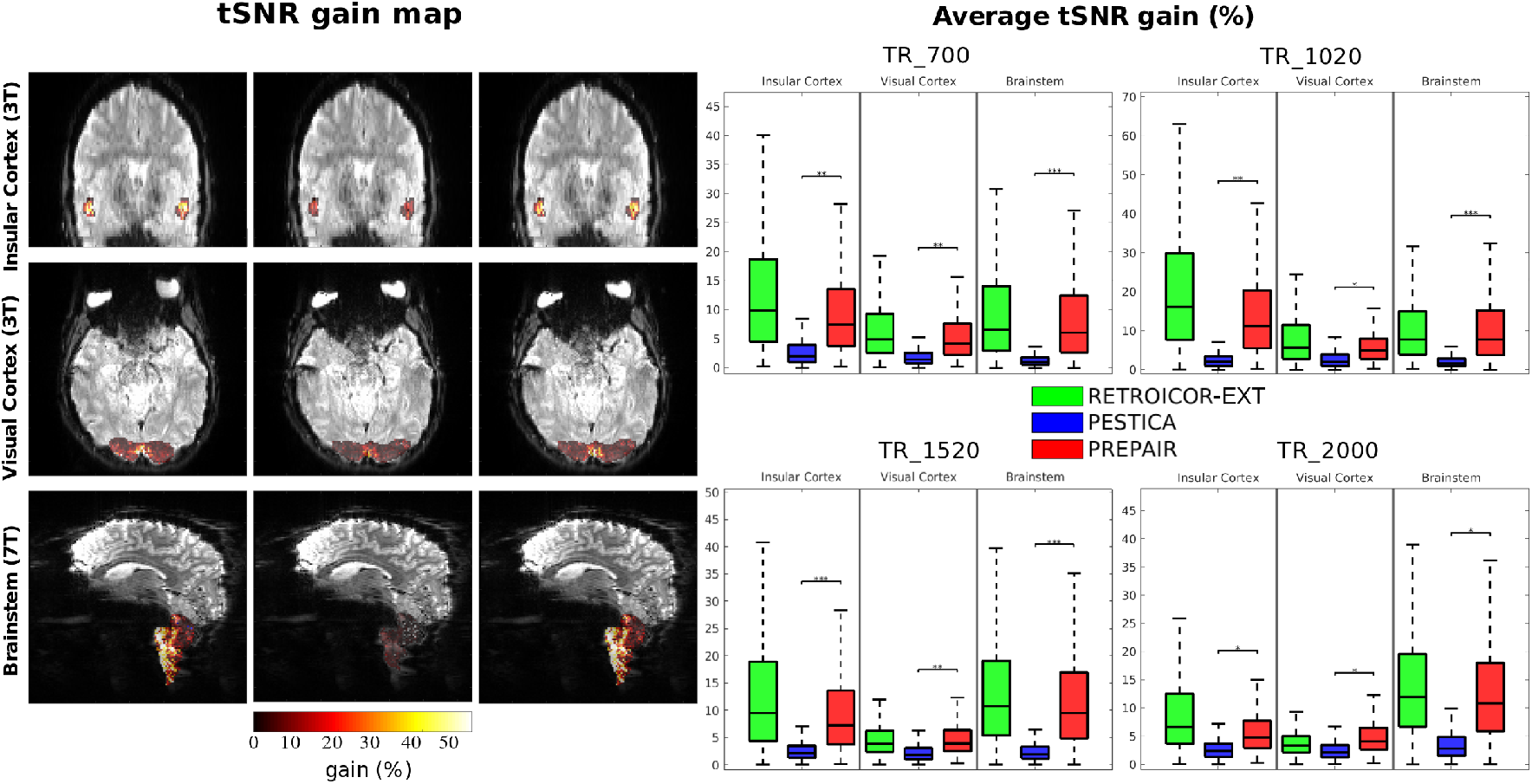
tSNR gain map (S2, 3T_TR_700) in different anatomical regions (left panel). The level of improvement with PREPAIR was comparable with RETROICOR-EXT and larger than with PESTICA. For all protocols and anatomical regions (right panel), tSNR gain with PREPAIR was significantly higher than PESTICA (*: p<0.05; **: p<0.01; ***: p<0.001). The level of tSNR improvement with PREPAIR in the insular and visual cortices is slightly weaker than RETROICOR-EXT but comparable in the brainstem.

We excluded FIX from this comparison, as we found a wide spread of standard deviations in the mean variance improvement across subjects (Supplementary Information sFig. 5), suggesting that other sources of noise and signal variation have been removed. Thus, tSNR improvements through FIX may not stem from the removal of the physiological noise, but from other signal and noise components. This was verified by identifying the FIX components that were not physiological-related, i.e. when one of the main three peaks of the power spectra did not correspond to the physiological fundamental frequency (or was aliased) with a tolerance of 10% around it. Over all protocols and subjects of the 3 T study, we found that 35% of the components selected by FIX were not physiological noise-related but related either to eye motion (Supplementary Information, sFig. 6) or to a resting state network (Supplementary Information, sFig. 7).

## 4. Discussion

We have developed a physiological data estimation and correction method called PREPAIR, which identifies physiological fluctuations from fMRI data without the need for supplementary external physiological recordings. The novel aspects of PREPAIR are that it uses both magnitude and phase information, physiological noise is sampled at the slice TR rather than the volume TR – so high frequency cardiac fluctuations are critically sampled – and it does not require user interaction to generate the magnitude or phase regressors or select which of the two best captures respiratory and cardiac fluctuations.

We attribute the utility of the phase in identifying respiration-related signals as being due to the sensitivity of the phase to change in the ΔB_0_ field where there is motion of structures with large susceptibility differences; primarily in this case the chest (Raj et al., 2001). Cardiac changes were generally better identified with the magnitude — 50% of the subjects and protocols at 3 T and 95% at 7 T (see Table 2) —, because of the large changes to the magnitude generated by spin history effects. In a few cases where the correlation between the phase and the external recordings was much higher than with the magnitude, PREPAIR would have failed if only the magnitude data had been used (see Supplementary Information sFig. 1).

Reliance on phase data could be viewed as a limitation of the PREPAIR approach, as phase images are not currently routinely reconstructed and stored in fMRI experiments. The use of phase requires a solution for the phase-sensitive combination of RF coil signals, but also phase unwrapping and the removal of unwanted signals from sources such as the cold-head helium pump which affect the phase signal to a larger extent than the magnitude fMRI signal (Hagberg et al., 2012). However, increasing interest in methods which use the phase for dynamic distortion correction (Dymerska et al., 2018), quantitative susceptibility mapping (QSM) (Sun and Wilman, 2015) and functional QSM (Balla et al., 2014) have led to the development of effective solutions to these problems.

Considering coil combination, MRI scanner vendors have effective coil combination methods for systems with a body coil (e.g. SENSE (Pruessmann et al., 1999) and pre-scan normalize combined with adaptive combination (Jellus and Kanengiesser, 2014)). For 7 T systems, a multi-echo ASPIRE pre-scan can be used to calculate phase offsets needed to combine separate channel EPI data (Eckstein et al., 2018) as was done here, with alternatives being offered by the CMRR sequence (Xu et al., 2013), which uses SENSE, and implementations of the VRC approach (Parker et al., 2014), to give a few examples.

For phase unwrapping, recent faster methods (Dymerska et al., 2021; Karsa and Shmueli, 2019) mean that this step can be performed sufficiently quickly and reliably that no longer poses an obstacle to phase imaging with EPI.

In regards to signals which disrupt the phase, the main frequency generated by the pump lies near the respiratory fundamental frequency for TR = 700 ms but without overlap, as seen in Figure 2. Our results have revealed that this perturbation would move away from the frequency range occupied by respiratory fluctuations with increasing TR, which agrees with Figure 1 in Hagberg et al. (2012) where the main cold pump peak lies at 0.67 Hz for a faster TR (180 ms), between typical respiration and cardiac frequency ranges. Protocols with TR below 700 ms would move this major perturbation to higher frequencies, which could limit the detection of the respiration fundamental frequency in the phase time series. Furthermore, the presence of sidebands can make it difficult to identify the fundamental physiological frequencies. Sidebands occur when a carrier signal is modulated by a lower frequency with high power, producing mirrored signals around the carrier. For instance, the coupling of the respiratory and the cardiac cycles – caused by the respiratory modulation of arterial-induced pulsations – leads to frequency contributions at the sum of and the difference between the fundamental cardiac frequency (the carrier) (Brooks et al., 2008; Raitamaa et al., 2021). The reordering process can also engender sidebands at f_n/TR_ ± f_m_ due to the amplitude modulation of the carrier signal f_n/TR_ (i.e. 1/TR and its harmonics) by a lower frequency f_m_, which can either be respiratory and helium cold pump frequencies in the case of phase data, or scanner drifts in the magnitude data. Those f_m_ frequencies are known to be dominant in fMRI data. This step can be critical for long TRs if the width of f_m_ is large (which is the case for respiratory signals but not for scanner drifts) as the resulting sidebands can lie much lower than typical cardiac frequencies and reach the respiratory domain. Because these only affects the phase signal, the magnitude time series will enable PREPAIR to overcome these challenges in either situation in case the phase fails to capture physiological signals. Indeed, for fast and low TRs, the magnitude component of PREPAIR proved to be able to detect respiratory and cardiac fluctuations with a very good agreement with the external recordings acquired at both 3 T and 7 T (see Table 2).

The slice signal detrending used slice TR sampling of physiological noise, but some volume TR dependency remains, and PREPAIR could fail to detect cardiac fundamental frequencies when they are too close to 1/TR frequency or harmonics thereof, especially for TR = 1020 ms (1/TR = 0.98 Hz) with subjects having a cardiac heart rate near 1 Hz, which is quite typical, or include the remains of this perturbation in the physiological waveforms. This could explain the poor correlation with the external physiological signals for TR_2000 (see Figure 3). Despite that, PREPAIR never significantly performed less well than RETROICOR-EXT in tSNR improvement and even performed comparably to RETROICOR-EXT in the brainstem (see Figure 6).

Our model did not include physiological noise originating in fluctuations in the heart rate (Akselrod et al., 1981; Cohen and Taylor, 2002; Otzenberger et al., 1998) and subtle changes in the depth and rate of breathing (Wise et al., 2004), which occur at much lower frequencies (<0.1 Hz) than that of the cardiac cycle (~1 Hz) and respiration (~0.3 Hz). Including these slower fluctuations as regressors in the analysis can improve the statistical power of fMRI studies and the detection of task-related BOLD signal changes (Birn et al., 2006; Chang et al., 2009; Shmueli et al., 2007); something which could be added to the PREPAIR output without any fundamental changes to the processing.

An important criterion for selecting a physiological noise correction method for multisubject studies is whether the approach can be executed without user intervention/unsupervised. FIX is challenging to use in a study with a large number of subjects. First, it is very time consuming, because MELODIC-ICA must be run prior to classification. Secondly, FIX is prone to include components of interest (i.e. signal changes related to activation) because of the dependence on the threshold parameter (see Supplementary Information sFig. 5), which controls the ratio of good to bad components, and which is subject dependent, as seen from the number of components labelled as physiological noise (see Section 2.5). This would explain the low performance of FIX at 7 T compared to 3 T. Finally but importantly, an optimal noise removal procedure with FIX requires manual checks that activation-related components are not excluded, which can be onerous. PESTICA can be run in either unsupervised or supervised mode. Running it in supervised mode involves defining the range of frequencies suspected to be physiological noise-related and may have improved its performance. The performance which could be achieved was assessed by inspecting all the power spectra of respiratory and cardiac estimators generated by PESTICA software (Supplementary Information sFig. 8 and sFig. 9; delta values are for the dispersion of the fundamental physiological frequencies with respect to those from the external recordings). Among these, narrowing the frequency distribution would have been challenging for 48 power spectra (60%) because either there was no obvious distribution (for instance Cardiac: S5_3T_TR_700, S9_3T_TR700; Respiration: S4_3T_TR_2000, S9_3T_TR_1520)), or there were multiple distributions (for instance Cardiac: S2_3T_TR_1520, S2_3T_TR_2000; Respiration: S9_3T_TR_1020, S10_3T_TR_2000), or there was a unique peak but this did not correlate with the expected fundamental physiological frequency (for instance Cardiac: S1_3T_TR_1520, S4_3T_TR_2000; Respiration: S3_3T_TR_1020, S9_3T_TR_700). Filtering the power spectra of 19 runs (23.75%) (for instance Cardiac: S2_3T_TR_700, S6_3T_TR_1020; Respiration: S2_3T_TR_1520, S7_3T_TR_1020) may have improved the quality of the estimators. Although the main peak was wrongly estimated, filtering around it would have included the expected value in the distribution. Finally, the quality of the power spectrum of 13 runs (16.25%) could likely have been improved because the fundamental physiological frequencies were correctly assessed (for instance Respiration: S1_3T_TR700, S6_3T_TR1020; Cardiac: S1_3T_TR_2000, S6_3T_TR_1520) and defining a narrower window, by hand, would have reduced the contribution of other non-physiological contributions to regressors (Respiration: S1_3T_TR_700 and S1_3T_TR_2000).

Finally, PREPAIR proved to be efficient in improving the variance in regions outside the cerebrum where imaging and the identification of the anatomical structure is challenging. The brainstem, for instance, is involved in disorders affecting autonomic disfunctions (Brook and Julius, 2000), affective disorders (Paul and Lowry, 2013), migraine (Denuelle and Fabre, 2013), and Parkinson’s disease (Braak et al., 2003; Holiga et al., 2015; Tison and Meissner, 2014), but is subject to high levels of physiological noise. The proximity of the brainstem to the fourth ventricle and arteries has made it the focus of effort to reduce the variance of contaminated voxels and improve sensitivity (Harvey et al., 2008; Beissner et al., 2011; Matt et al., 2019) to allow the depiction of quite small nuclei (D’Ardenne et al., 2008; Thompson et al., 2006). This can be achieved without the need for external physiological recordings using an anatomically-defined mask of brainstem and ICA (Beissner et al., 2014). Combining this with temporal noise regression would reduce variance further, provided that the physiological noise removal tools did not remove signals of interests or add noise to the data, which would impede the detection of neurally-driven responses, as shown by Agrawal et al. (2020) with the method CompCor (Behzadi et al., 2007). Our results have demonstrated that PREPAIR performed similarly to RETROICOR-EXT and PESTICA, and significantly better than FIX, in the preservation of the integrity of the magnitude signal outside areas affected by physiological noise in the 3 T and 7 T studies (see Table 3 and Table 4).

## Conclusions

The proposed physiological noise correction method PREPAIR uses time series phase and magnitude images to identify respiratory and cardiac fluctuations in fMRI, unsupervised and without external recordings. The physiological signals identified by PREPAIR agreed well with those from the respiratory bellows and photoplethysmography and led to a similar quality of correction to a method based on external recordings – RETROICOR-EXT – and ICA-based FIX at 3 T, and better than other recording-free noise correction method PESTICA, at both 3 T and 7 T.

## Acknowledgements

This study was funded by the Austrian Science Fund (FWF) project 31452. SR was additionally supported by the Marie Skłodowska-Curie Action (MS-fMRI-QSM 794298).

## References

Agrawal, U., Brown, E.N., Lewis, L.D., 2020. Model-based physiological noise removal in fast fMRI. Neuroimage 205, 116231. https://doi.org/10.1016/j.neuroimage.2019.116231

Akselrod, S., Gordon, D., Ubel, F.A., Shannon, D.C., Berger, A.C., Cohen, R.J., 1981. Power spectrum analysis of heart rate fluctuation: a quantitative probe of beat-to-beat cardiovascular control. Science 213, 220–222. https://doi.org/10.1126/science.6166045

Aslan, S., Hocke, L., Schwarz, N., Frederick, B., 2019. Extraction of the cardiac waveform from simultaneous multislice fMRI data using slice sorted averaging and a deep learning reconstruction filter. Neuroimage 198, 303–316. https://doi.org/10.1016/j.neuroimage.2019.05.049

Bachrata, B., Eckstein, K., Trattnig, S., Robinson, S.D., 2018. Considerations in quantitative susceptibility mapping using echo-planar imaging. Proc. ISMRM, Joint Annual Meeting, Paris, France #4996.

Balla, D.Z., Sanchez-Panchuelo, R.M., Wharton, S.J., Hagberg, G.E., Scheffler, K., Francis, S.T., Bowtell, R., 2014. Functional quantitative susceptibility mapping (fQSM). Neuroimage 100, 112–124. https://doi.org/10.1016/j.neuroimage.2014.06.011

Barth, M., Breuer, F., Koopmans, P.J., Norris, D.G., Poser, B.A., 2016. Simultaneous multislice (SMS) imaging techniques. Magn Reson Med 75, 63–81. https://doi.org/10.1002/mrm.25897

Beall, E.B., 2010. Adaptive cyclic physiologic noise modeling and correction in functional MRI. J Neurosci Methods 187, 216–228. https://doi.org/10.1016/j.jneumeth.2010.01.013

Beall, E.B., Lowe, M.J., 2007. Isolating physiologic noise sources with independently determined spatial measures. Neuroimage 37, 1286–1300. https://doi.org/10.1016/j.neuroimage.2007.07.004

Behzadi, Y., Restom, K., Liau, J., Liu, T.T., 2007. A component based noise correction method (CompCor) for BOLD and perfusion based fMRI. Neuroimage 37, 90–101. https://doi.org/10.1016/j.neuroimage.2007.04.042

Beissner, F., Deichmann, R., Baudrexel, S., 2011. fMRI of the brainstem using dual-echo EPI. NeuroImage 55, 1593–1599. https://doi.org/10.1016/j.neuroimage.2011.01.042

Beissner, F., Schumann, A., Brunn, F., Eisenträger, D., Bär, K.-J., 2014. Advances in functional magnetic resonance imaging of the human brainstem. NeuroImage 86, 91–98. https://doi.org/10.1016/j.neuroimage.2013.07.081

Bianciardi, M., Fukunaga, M., van Gelderen, P., Horovitz, S.G., de Zwart, J.A., Shmueli, K., Duyn, J.H., 2009. Sources of fMRI signal fluctuations in the human brain at rest: a 7T study. Magn Reson Imaging 27, 1019–1029. https://doi.org/10.1016/j.mri.2009.02.004

Birn, R.M., Diamond, J.B., Smith, M.A., Bandettini, P.A., 2006. Separating respiratory-variation-related fluctuations from neuronal-activity-related fluctuations in fMRI. Neuroimage 31, 1536–1548. https://doi.org/10.1016/j.neuroimage.2006.02.048

Birn, R.M., Murphy, K., Bandettini, P.A., 2008. The effect of respiration variations on independent component analysis results of resting state functional connectivity. Hum Brain Mapp 29, 740–750. https://doi.org/10.1002/hbm.20577

Biswal, B., DeYoe, E.A., Hyde, J.S., 1996. Reduction of physiological fluctuations in fMRI using digital filters. Magn Reson Med 35, 107–113. https://doi.org/10.1002/mrm.1910350114

Bokil, H., Andrews, P., Kulkarni, J.E., Mehta, S., Mitra, P.P., 2010. Chronux: a platform for analyzing neural signals. J Neurosci Methods 192, 146–151. https://doi.org/10.1016/j.jneumeth.2010.06.020

Braak, H., Tredici, K.D., Rüb, U., de Vos, R.A.I., Jansen Steur, E.N.H., Braak, E., 2003. Staging of brain pathology related to sporadic Parkinson’s disease. Neurobiology of Aging 24, 197–211. https://doi.org/10.1016/S0197-4580(02)00065-9

Brook, R.D., Julius, S., 2000. Autonomic imbalance, hypertension, and cardiovascular risk. American Journal of Hypertension 13, 112S–122S. https://doi.org/10.1016/S0895-7061(00)00228-4

Brooks, J.C.W., Beckmann, C.F., Miller, K.L., Wise, R.G., Porro, C.A., Tracey, I., Jenkinson, M., 2008. Physiological noise modelling for spinal functional magnetic resonance imaging studies. Neuroimage 39, 680–692. https://doi.org/10.1016/j.neuroimage.2007.09.018

Caballero-Gaudes, C., Reynolds, R.C., 2017. Methods for cleaning the BOLD fMRI signal. Neuroimage 154, 128–149. https://doi.org/10.1016/j.neuroimage.2016.12.018

Chang, C., Cunningham, J.P., Glover, G.H., 2009. Influence of heart rate on the BOLD signal: the cardiac response function. Neuroimage 44, 857–869. https://doi.org/10.1016/j.neuroimage.2008.09.029

Cheng, H., Li, Y., 2010. Respiratory noise correction using phase information. Magnetic Resonance Imaging 28, 574–582. https://doi.org/10.1016/j.mri.2009.12.014

Chuang, K.H., Chen, J.H., 2001. IMPACT: image-based physiological artifacts estimation and correction technique for functional MRI. Magn Reson Med 46, 344–353. https://doi.org/10.1002/mrm.1197

Churchill, N.W., Strother, S.C., 2013. PHYCAA+: An optimized, adaptive procedure for measuring and controlling physiological noise in BOLD fMRI. NeuroImage 82, 306–325. https://doi.org/10.1016/j.neuroimage.2013.05.102

Cohen, M.A., Taylor, J.A., 2002. Short-term cardiovascular oscillations in man: measuring and modelling the physiologies. J Physiol 542, 669–683. https://doi.org/10.1113/jphysiol.2002.017483

Cox, R.W., 1996. AFNI: software for analysis and visualization of functional magnetic resonance neuroimages. Comput Biomed Res 29, 162–173. https://doi.org/10.1006/cbmr.1996.0014

Cox, R.W., Hyde, J.S., 1997. Software tools for analysis and visualization of fMRI data. NMR Biomed 10, 171–178. https://doi.org/10.1002/(sici)1099-1492(199706/08)10:4/5<171::aid-nbm453>3.0.co;2-l

Curtis, A.T., Menon, R.S., 2014. Highcor: A novel data-driven regressor identification method for BOLD fMRI. NeuroImage 98, 184–194. https://doi.org/10.1016/j.neuroimage.2014.05.013

Dagli, M.S., Ingeholm, J.E., Haxby, J.V., 1999. Localization of cardiac-induced signal change in fMRI. Neuroimage 9, 407–415. https://doi.org/10.1006/nimg.1998.0424

D’Ardenne, K., McClure, S.M., Nystrom, L.E., Cohen, J.D., 2008. BOLD Responses Reflecting Dopaminergic Signals in the Human Ventral Tegmental Area. Science 319, 1264–1267. https://doi.org/10.1126/science.1150605

Denuelle, M., Fabre, N., 2013. Functional neuroimaging of migraine. Revue Neurologique, Migraine 169, 380–389. https://doi.org/10.1016/j.neurol.2013.02.002

Dymerska, B., Eckstein, K., Bachrata, B., Siow, B., Trattnig, S., Shmueli, K., Robinson, S.D., 2021. Phase unwrapping with a rapid opensource minimum spanning tree algorithm (ROMEO). Magn Reson Med 85, 2294–2308. https://doi.org/10.1002/mrm.28563

Dymerska, B., Poser, B.A., Barth, M., Trattnig, S., Robinson, S.D., 2018. A method for the dynamic correction of B0-related distortions in single-echo EPI at 7T. Neuroimage 168, 321–331. https://doi.org/10.1016/j.neuroimage.2016.07.009

Eckstein, K., Dymerska, B., Bachrata, B., Bogner, W., Poljanc, K., Trattnig, S., Robinson, S.D., 2018. Computationally Efficient Combination of Multi-channel Phase Data From Multi-echo Acquisitions (ASPIRE). Magn Reson Med 79, 2996–3006. https://doi.org/10.1002/mrm.26963

Elgendi, M., 2012. On the Analysis of Fingertip Photoplethysmogram Signals. Current Cardiology Reviews 8, 14–25.

Feinberg, D.A., Moeller, S., Smith, S.M., Auerbach, E., Ramanna, S., Gunther, M., Glasser, M.F., Miller, K.L., Ugurbil, K., Yacoub, E., 2010. Multiplexed echo planar imaging for sub-second whole brain FMRI and fast diffusion imaging. PLoS One 5, e15710. https://doi.org/10.1371/journal.pone.0015710

Feinberg, D.A., Setsompop, K., 2013. Ultra-fast MRI of the human brain with simultaneous multi-slice imaging. J Magn Reson 229, 90–100. https://doi.org/10.1016/j.jmr.2013.02.002

Frank, L.R., Buxton, R.B., Wong, E.C., 2001. Estimation of respiration-induced noise fluctuations from undersampled multislice fMRI data. Magn Reson Med 45, 635–644. https://doi.org/10.1002/mrm.1086

Glover, G.H., Li, T.Q., Ress, D., 2000. Image-based method for retrospective correction of physiological motion effects in fMRI: RETROICOR. Magn Reson Med 44, 162–167. https://doi.org/10.1002/1522-2594(200007)44:1<162::aid-mrm23>3.0.co;2-e

Hagberg, G.E., Bianciardi, M., Brainovich, V., Cassara, A.M., Maraviglia, B., 2012. Phase stability in fMRI time series: effect of noise regression, off-resonance correction and spatial filtering techniques. Neuroimage 59, 3748–3761. https://doi.org/10.1016/j.neuroimage.2011.10.095

Hagberg, G.E., Bianciardi, M., Brainovich, V., Cassarà, A.M., Maraviglia, B., 2008. The effect of physiological noise in phase functional magnetic resonance imaging: from blood oxygen level-dependent effects to direct detection of neuronal currents. Magn Reson Imaging 26, 1026–1040. https://doi.org/10.1016/j.mri.2008.01.010

Harvey, A.K., Pattinson, K.T.S., Brooks, J.C.W., Mayhew, S.D., Jenkinson, M., Wise, R.G., 2008. Brainstem functional magnetic resonance imaging: disentangling signal from physiological noise. J Magn Reson Imaging 28, 1337–1344. https://doi.org/10.1002/jmri.21623

Holiga, Š., Mueller, K., Möller, H.E., Urgošík, D., Růžička, E., Schroeter, M.L., Jech, R., 2015. Resting-state functional magnetic resonance imaging of the subthalamic microlesion and stimulation effects in Parkinson’s disease: Indications of a principal role of the brainstem. NeuroImage: Clinical 9, 264–274. https://doi.org/10.1016/j.nicl.2015.08.008

Jellus, V., Kanengiesser, V., 2014. Adaptive Coil Combination Using a Body Coil Scan as Phase Reference. Proc. ISMRM, 23th Joint Annual Meeting, Paris, France #4406.

Jezzard, P., LeBihan, D., Cuenod, C., Pannier, L., Prinster, A., Turner, R., 1993. An Investigation of the Contribution of Physiological Noise in Human Functional MRI Studies at 1.5 Tesla and 4 Tesla. Proc. ISMRM, 12th Annual Meeting, New York, USA #1993, 1392.

Karsa, A., Shmueli, K., 2019. SEGUE: A Speedy rEgion-Growing Algorithm for Unwrapping Estimated Phase. IEEE Trans Med Imaging 38, 1347–1357. https://doi.org/10.1109/TMI.2018.2884093

Kasper, L., Bollmann, S., Diaconescu, A.O., Hutton, C., Heinzle, J., Iglesias, S., Hauser, T.U., Sebold, M., Manjaly, Z.-M., Pruessmann, K.P., Stephan, K.E., 2017. The PhysIO Toolbox for Modeling Physiological Noise in fMRI Data. J Neurosci Methods 276, 56–72. https://doi.org/10.1016/j.jneumeth.2016.10.019

Krüger, G., Glover, G.H., 2001. Physiological noise in oxygenation-sensitive magnetic resonance imaging. Magn Reson Med 46, 631–637. https://doi.org/10.1002/mrm.1240

Le, T.H., Hu, X., 1996. Retrospective estimation and correction of physiological artifacts in fMRI by direct extraction of physiological activity from MR data. Magn Reson Med 35, 290–298. https://doi.org/10.1002/mrm.1910350305

Liu, T.T., 2016. Noise contributions to the fMRI signal: An overview. Neuroimage 143, 141–151. https://doi.org/10.1016/j.neuroimage.2016.09.008

Lund, T.E., Madsen, K.H., Sidaros, K., Luo, W.-L., Nichols, T.E., 2006. Non-white noise in fMRI: does modelling have an impact? Neuroimage 29, 54–66. https://doi.org/10.1016/j.neuroimage.2005.07.005

Matt, E., Fischmeister, F.P.S., Amini, A., Robinson, S.D., Weber, A., Foki, T., Gizewski, E.R., Beisteiner, R., 2019. Improving sensitivity, specificity, and reproducibility of individual brainstem activation. Brain Struct Funct 224, 2823–2838. https://doi.org/10.1007/s00429-019-01936-3

Ogawa, S., Lee, T.M., Kay, A.R., Tank, D.W., 1990. Brain magnetic resonance imaging with contrast dependent on blood oxygenation. Proc Natl Acad Sci U S A 87, 9868–9872.

Otzenberger, H., Gronfier, C., Simon, C., Charloux, A., Ehrhart, J., Piquard, F., Brandenberger, G., 1998. Dynamic heart rate variability: a tool for exploring sympathovagal balance continuously during sleep in men. Am J Physiol 275, H946–950. https://doi.org/10.1152/ajpheart.1998.275.3.H946

Parker, D.L., Payne, A., Todd, N., Hadley, J.R., 2014. Phase reconstruction from multiple coil data using a virtual reference coil. Magnetic Resonance in Medicine 72, 563–569. https://doi.org/10.1002/mrm.24932

Paul, E.D., Lowry, C.A., 2013. Functional topography of serotonergic systems supports the Deakin/Graeff hypothesis of anxiety and affective disorders. J Psychopharmacol 27, 1090–1106. https://doi.org/10.1177/0269881113490328

Perlbarg, V., Bellec, P., Anton, J.-L., Pélégrini-Issac, M., Doyon, J., Benali, H., 2007. CORSICA: correction of structured noise in fMRI by automatic identification of ICA components. Magn Reson Imaging 25, 35–46. https://doi.org/10.1016/j.mri.2006.09.042

Petridou, N., Schäfer, A., Gowland, P., Bowtell, R., 2009. Phase vs. magnitude information in functional magnetic resonance imaging time series: toward understanding the noise. Magn Reson Imaging 27, 1046–1057. https://doi.org/10.1016/j.mri.2009.02.006

Pruessmann, K.P., Weiger, M., Scheidegger, M.B., Boesiger, P., 1999. SENSE: sensitivity encoding for fast MRI. Magn Reson Med 42, 952–962.

Raitamaa, L., Huotari, N., Korhonen, V., Helakari, H., Koivula, A., Kananen, J., Kiviniemi, V., 2021. Spectral analysis of physiological brain pulsations affecting the BOLD signal. Human Brain Mapping 42, 4298–4313. https://doi.org/10.1002/hbm.25547

Raj, D., Anderson, A.W., Gore, J.C., 2001. Respiratory effects in human functional magnetic resonance imaging due to bulk susceptibility changes. Phys Med Biol 46, 3331–3340. https://doi.org/10.1088/0031-9155/46/12/318

Reynaud, O., Jorge, J., Gruetter, R., Marques, J.P., van der Zwaag, W., 2017. Influence of physiological noise on accelerated 2D and 3D resting state functional MRI data at 7 T. Magn Reson Med 78, 888–896. https://doi.org/10.1002/mrm.26823

Robinson, S.D., Bachrata, B., Eckstein, K., Trattnig, S., Enzinger, C., Barth, M., 2021. Improved dynamic distortion correction for fMRI using single-echo EPI, a fast sensivity scan and readout-reversed first image (REFILL). Proceedings of the 2021 ISMRM & SMRT Annual Meeting & Exhibition (Virtual) 671.

Robinson, S.D., Bredies, K., Khabipova, D., Dymerska, B., Marques, J.P., Schweser, F., 2017. An illustrated comparison of processing methods for MR phase imaging and QSM: combining array coil signals and phase unwrapping. NMR in Biomedicine 30, e3601. https://doi.org/10.1002/nbm.3601

Rowe, D.B., 2005. Modeling both the magnitude and phase of complex-valued fMRI data. Neuroimage 25, 1310–1324. https://doi.org/10.1016/j.neuroimage.2005.01.034

Salimi-Khorshidi, G., Douaud, G., Beckmann, C.F., Glasser, M.F., Griffanti, L., Smith, S.M., 2014. Automatic denoising of functional MRI data: Combining independent component analysis and hierarchical fusion of classifiers. NeuroImage 90, 449–468. https://doi.org/10.1016/j.neuroimage.2013.11.046

Setsompop, K., Gagoski, B.A., Polimeni, J.R., Witzel, T., Wedeen, V.J., Wald, L.L., 2012. Blipped-controlled aliasing in parallel imaging for simultaneous multislice echo planar imaging with reduced g-factor penalty. Magn Reson Med 67, 1210–1224. https://doi.org/10.1002/mrm.23097

Shin, W., Beall, E.B., Lowe, M.J., 2016. PESTICA 3.0: Evaluation of a new Physiologic estimation by temporal indepedent components analysis. Proc. OHBM 22nd Annual Meeting #4318.

Shmueli, K., van Gelderen, P., de Zwart, J.A., Horovitz, S.G., Fukunaga, M., Jansma, J.M., Duyn, J.H., 2007. Low-frequency fluctuations in the cardiac rate as a source of variance in the resting-state fMRI BOLD signal. Neuroimage 38, 306–320. https://doi.org/10.1016/j.neuroimage.2007.07.037

Sun, H., Wilman, A.H., 2015. Quantitative susceptibility mapping using single-shot echo-planar imaging. Magn Reson Med 73, 1932–1938. https://doi.org/10.1002/mrm.25316

Thomas, C.G., Harshman, R.A., Menon, R.S., 2002. Noise Reduction in BOLD-Based fMRI Using Component Analysis. NeuroImage 17, 1521–1537. https://doi.org/10.1006/nimg.2002.1200

Thompson, S.K., von Kriegstein, K., Deane-Pratt, A., Marquardt, T., Deichmann, R., Griffiths, T.D., McAlpine, D., 2006. Representation of interaural time delay in the human auditory midbrain. Nat Neurosci 9, 1096–1098. https://doi.org/10.1038/nn1755

Tison, F., Meissner, W.G., 2014. Movement disorders in 2013: diagnosing and treating PD-the earlier the better? Nat Rev Neurol 10, 65–66. https://doi.org/10.1038/nrneurol.2013.273

Triantafyllou, C., Hoge, R.D., Krueger, G., Wiggins, C.J., Potthast, A., Wiggins, G.C., Wald, L.L., 2005. Comparison of physiological noise at 1.5 T, 3 T and 7 T and optimization of fMRI acquisition parameters. Neuroimage 26, 243–250. https://doi.org/10.1016/j.neuroimage.2005.01.007

Vizioli, L., Moeller, S., Dowdle, L., Akçakaya, M., De Martino, F., Yacoub, E., Uğurbil, K., 2021. Lowering the thermal noise barrier in functional brain mapping with magnetic resonance imaging. Nat Commun 12, 5181. https://doi.org/10.1038/s41467-021-25431-8

Weisskoff, R., Baker, J., Belliveau, J., Davis, T., Kwong, K., Cohen, M., Rosen, B., 1993. Power Spectrum Analysis of Functionally-Weighted MR Data: What’s in the noise? Proc. ISMRM, 12th Annual Meeting, New York, USA #1993, 7.

Windischberger, C., Langenberger, H., Sycha, T., Tschernko, E.M., Fuchsjäger-Mayerl, G., Schmetterer, L., Moser, E., 2002. On the origin of respiratory artifacts in BOLD-EPI of the human brain. Magn Reson Imaging 20, 575–582. https://doi.org/10.1016/s0730-725x(02)00563-5

Wise, R.G., Ide, K., Poulin, M.J., Tracey, I., 2004. Resting fluctuations in arterial carbon dioxide induce significant low frequency variations in BOLD signal. Neuroimage 21, 1652–1664. https://doi.org/10.1016/j.neuroimage.2003.11.025

Xu, J., Moeller, S., Auerbach, E.J., Strupp, J., Smith, S.M., Feinberg, D.A., Yacoub, E., Uğurbil, K., 2013. Evaluation of slice accelerations using multiband echo planar imaging at 3 T. Neuroimage 83, 991–1001. https://doi.org/10.1016/j.neuroimage.2013.07.055

Zahneisen, B., Assländer, J., LeVan, P., Hugger, T., Reisert, M., Ernst, T., Hennig, J., 2014. Quantification and correction of respiration induced dynamic field map changes in fMRI using 3D single shot techniques. Magn Reson Med 71, 1093–1102. https://doi.org/10.1002/mrm.24771

