## Supplementary Material for "Unsupervised physiological noise correction of fMRI data using phase and magnitude information (PREPAIR)"

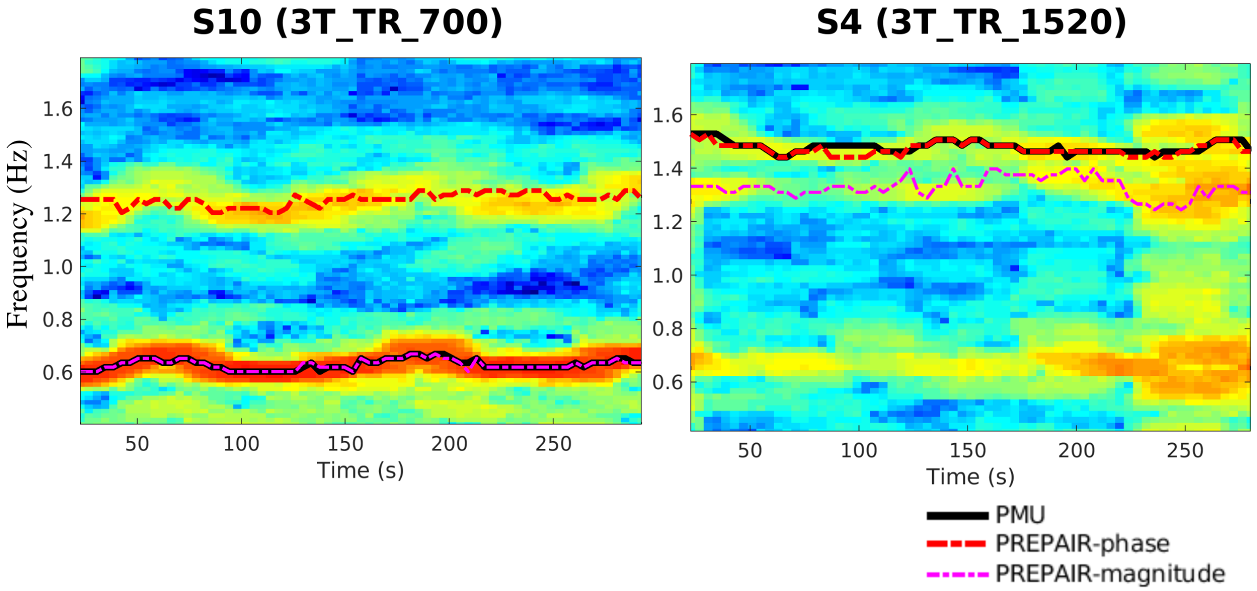


*sFig. 1: Spectrograms of two subjects with low (left: S10 with a cardiac frequency of ~0.6 Hz) and high (right: S4 with a cardiac frequency of ~1.5 Hz) cardiac dynamics, identified only by PREPAIR-magnitude (magenta) in the first case and only PREPAIR-phase(blue) in the second case. In both cases, the PREPAIR cardiac frequencies estimates overlay well with those identified by the PMU (black).*

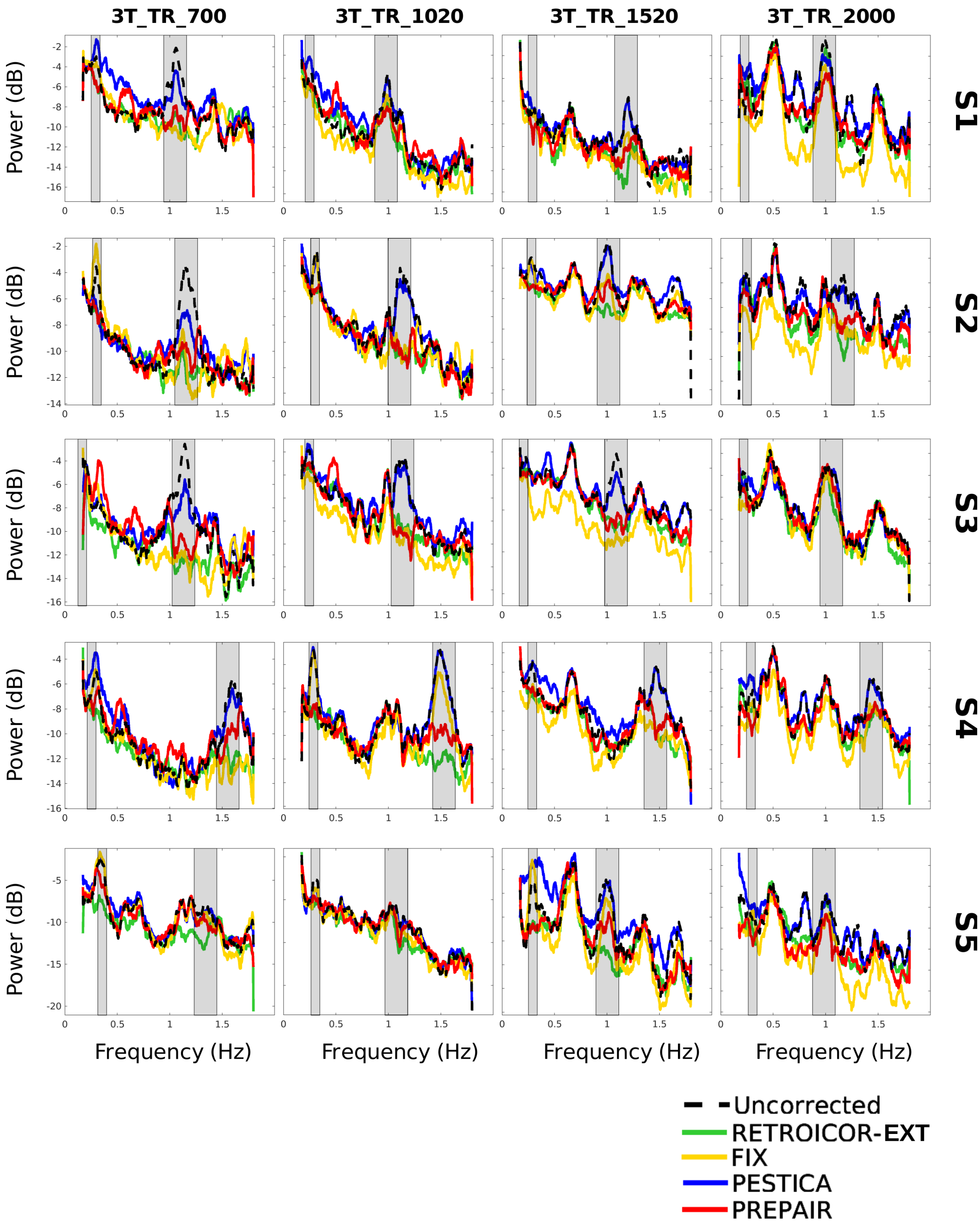


*sFig. 2: Comparison of the power spectrum of the uncorrected (dashed black line) and corrected (green, yellow, blue, and red lines for RETROICOR, FIX, PESTICA and PREPAIR, respectively) magnitude data for the first five subjects of the 3 T whole brain study (rows) for each protocol (columns). Grey boxes are for the location of the 1st harmonic (and 2nd when applicable) of respiratory and cardiac noise. For a better visualization, all frequency distribution were smoothed.*

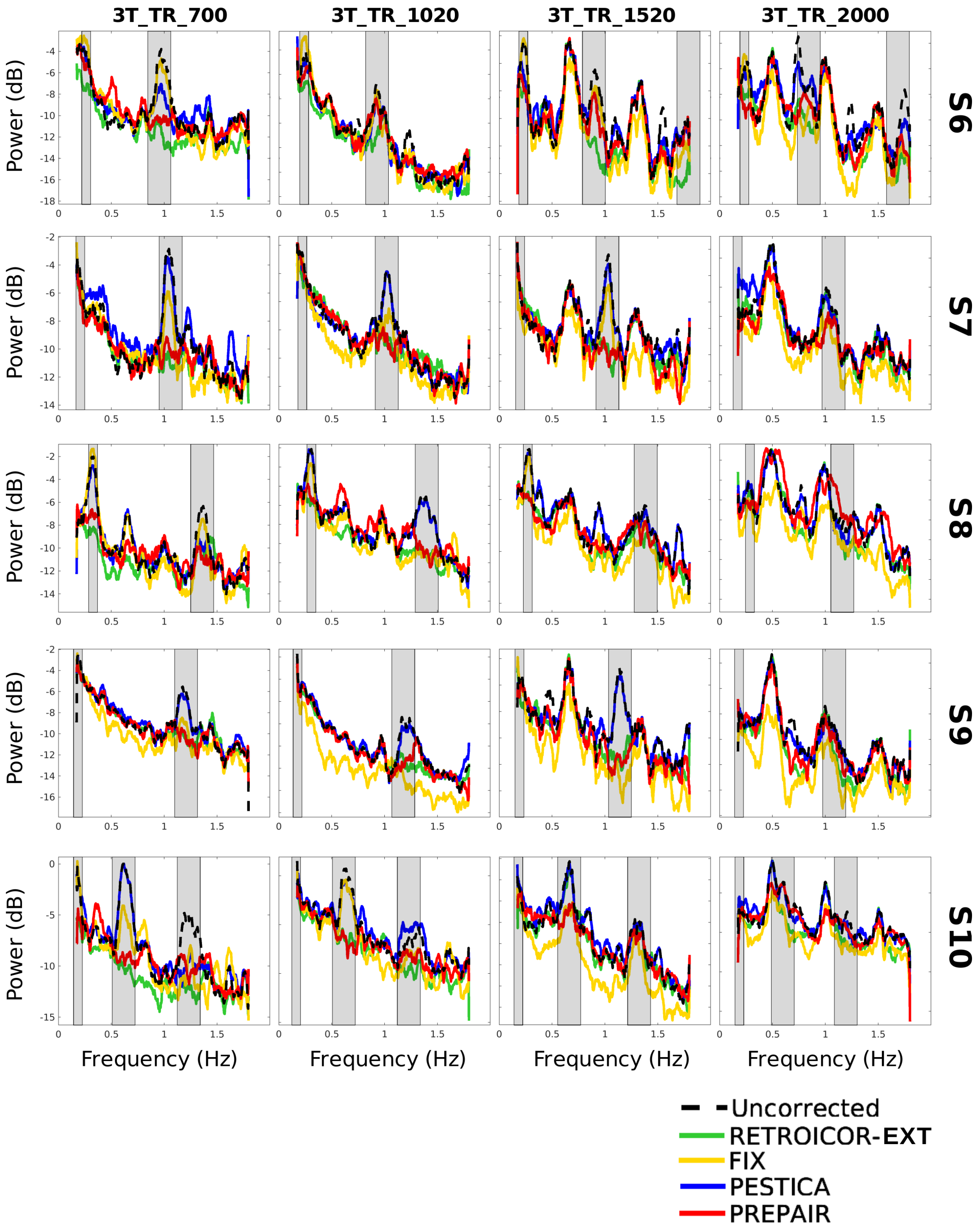


*sFig. 3: Comparison of the power spectrum of the uncorrected (dashed black line) and corrected (green, yellow, blue, and red lines for RETROICOR, FIX, PESTICA and PREPAIR, respectively) magnitude data for the last five subjects of the 3 T whole brain study (rows) for each protocol (columns). Grey boxes are for the location of the 1st harmonic (and 2nd when applicable) of respiratory and cardiac noise. For a better visualization, all frequency distribution were smoothed.*

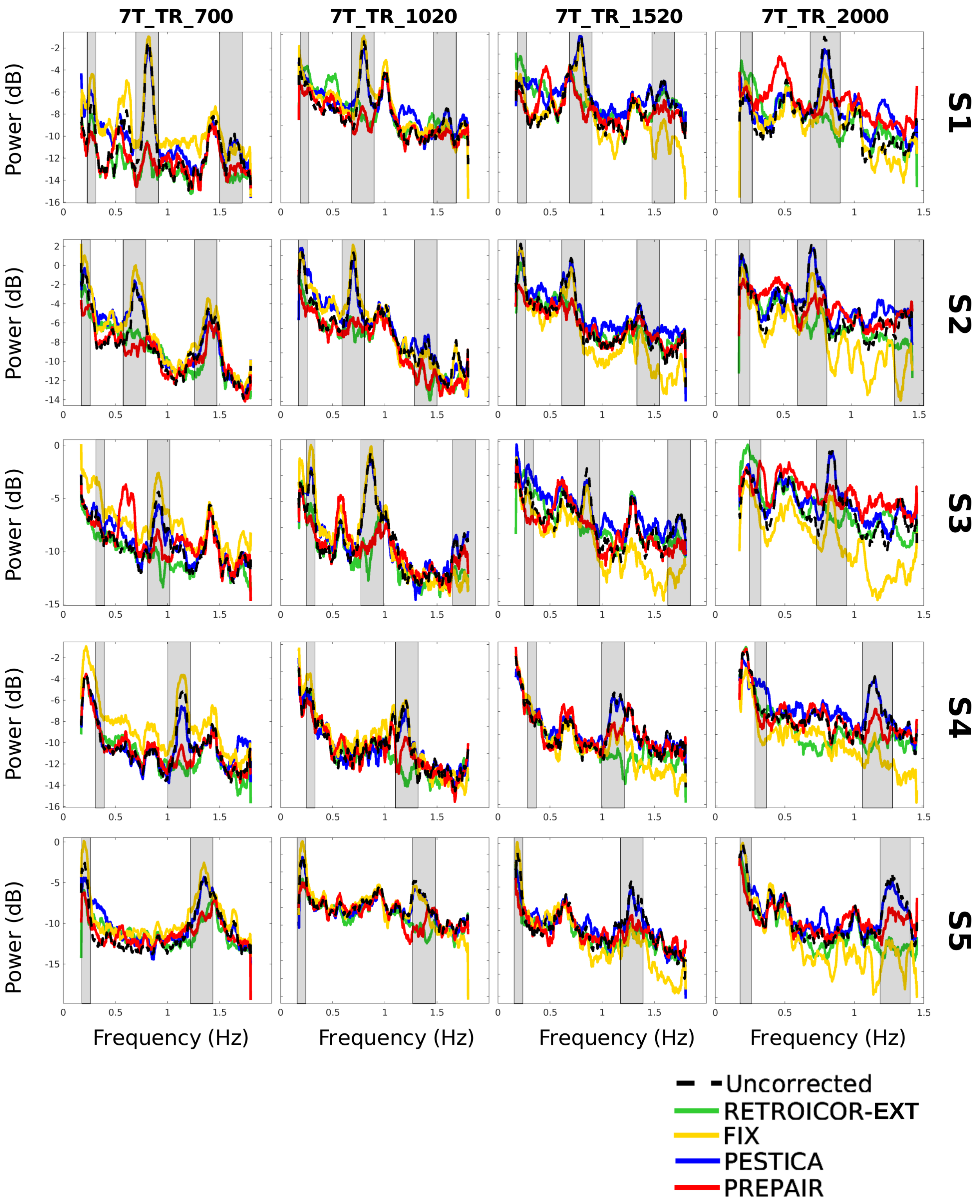


*sFig. 4: Comparison of the power spectrum of the uncorrected (dashed black line) and corrected (green, yellow, blue, and red lines for RETROICOR, FIX, PESTICA and PREPAIR, respectively) magnitude data for 7 T brainstem study (rows) for each protocol (columns). Grey boxes are for the location of the 1st harmonic (and 2nd when applicable) of respiratory and cardiac noise. For a better visualization, all frequency distribution were smoothed.*

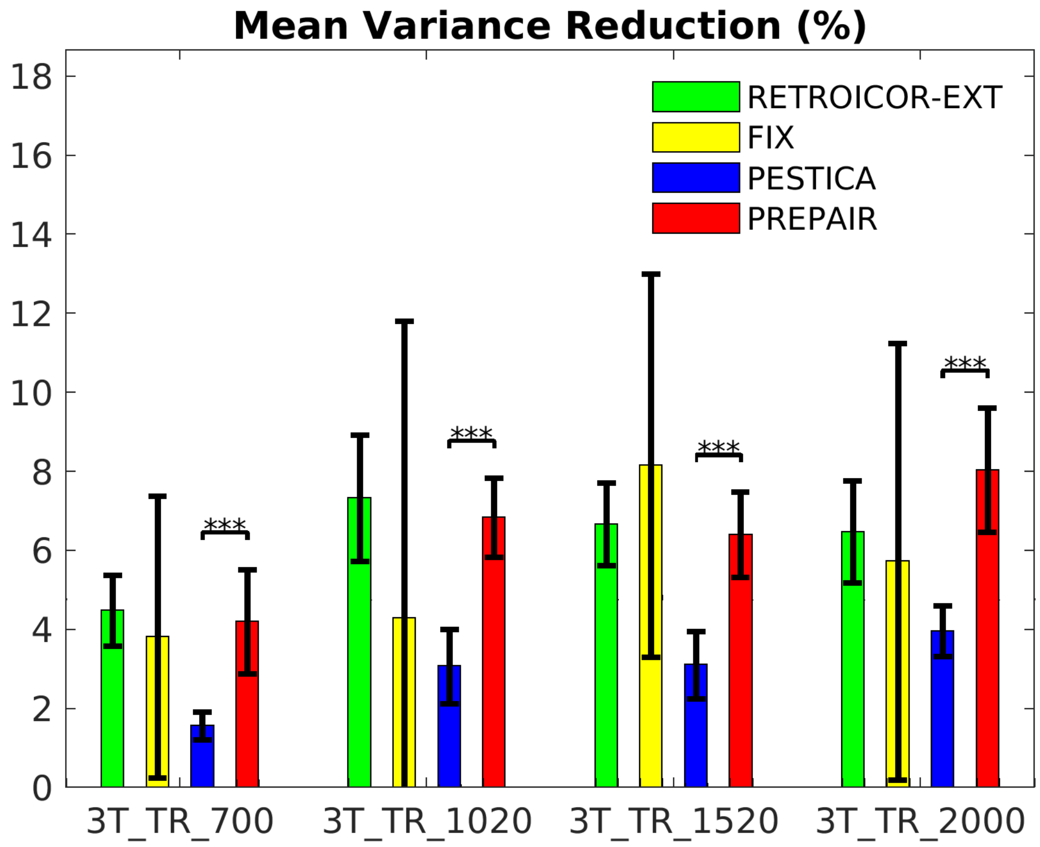


*sFig. 5: Variance improvement over all subjects. Because of additional unrelated physiological noise removed in some subjects, FIX would outperform PREPAIR. For all protocols, PREPAIR performed similarly as RETROICOR-EXT and significantly better than PESTICA (***: p<0.001).*

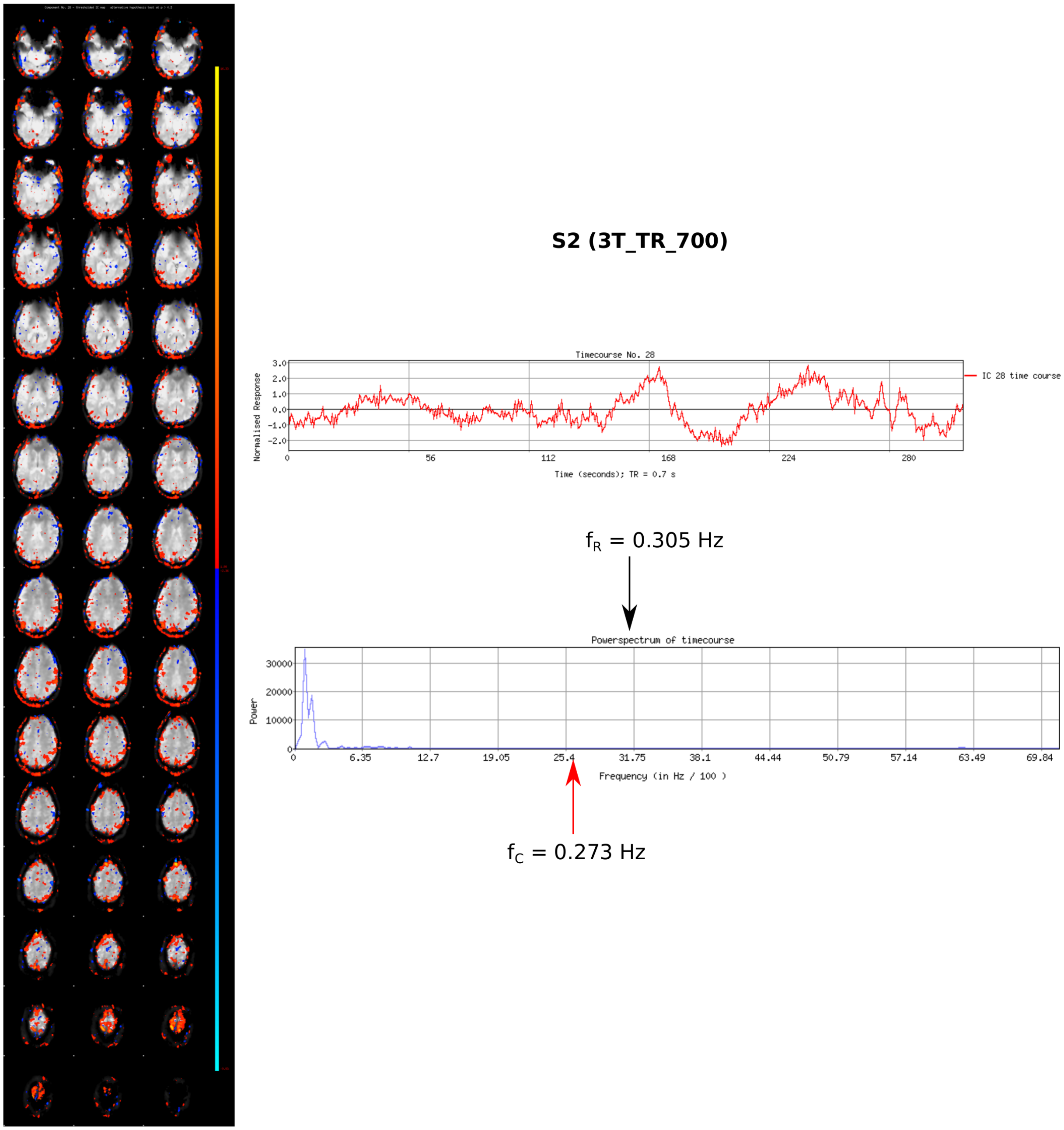


*sFig. 6: Eye motion-dominated component identified and as physiological noise by FIX: an example contribution to the wide spread of standard deviations in ssFig. 5. Fundamental cardiac $f_C$ (aliases) and respiratory $f_R$ derived from the PMU are shown on the power spectra.*

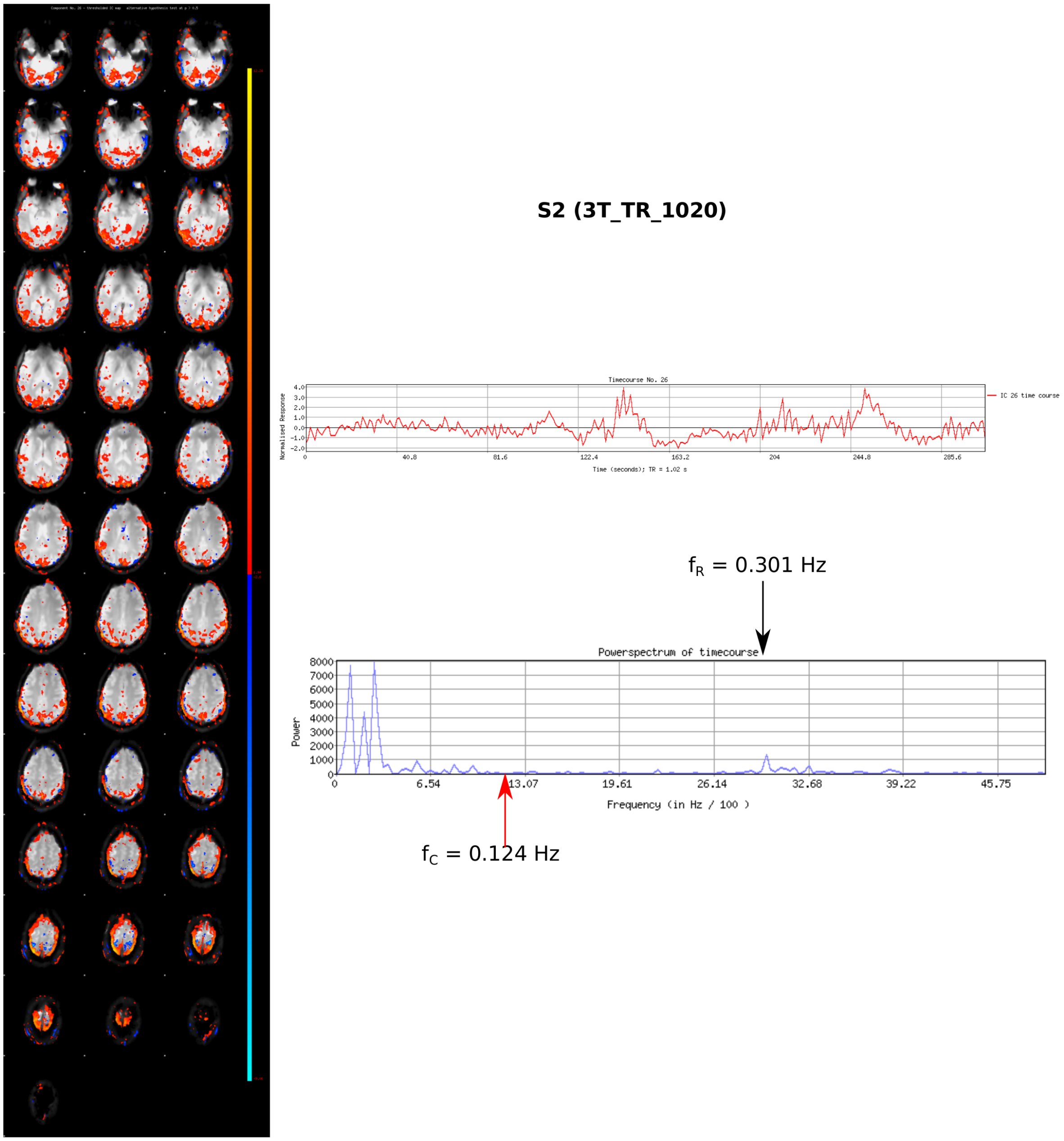


*sFig. 7: Resting state network-and head motion-related component identified as physiological noise by FIX: an example contribution to the wide spread of standard deviations in ssFig. 5. Fundamental cardiac $f_C$ (aliases) and respiratory $f_R$ derived from the PMU are shown on the power spectra.*

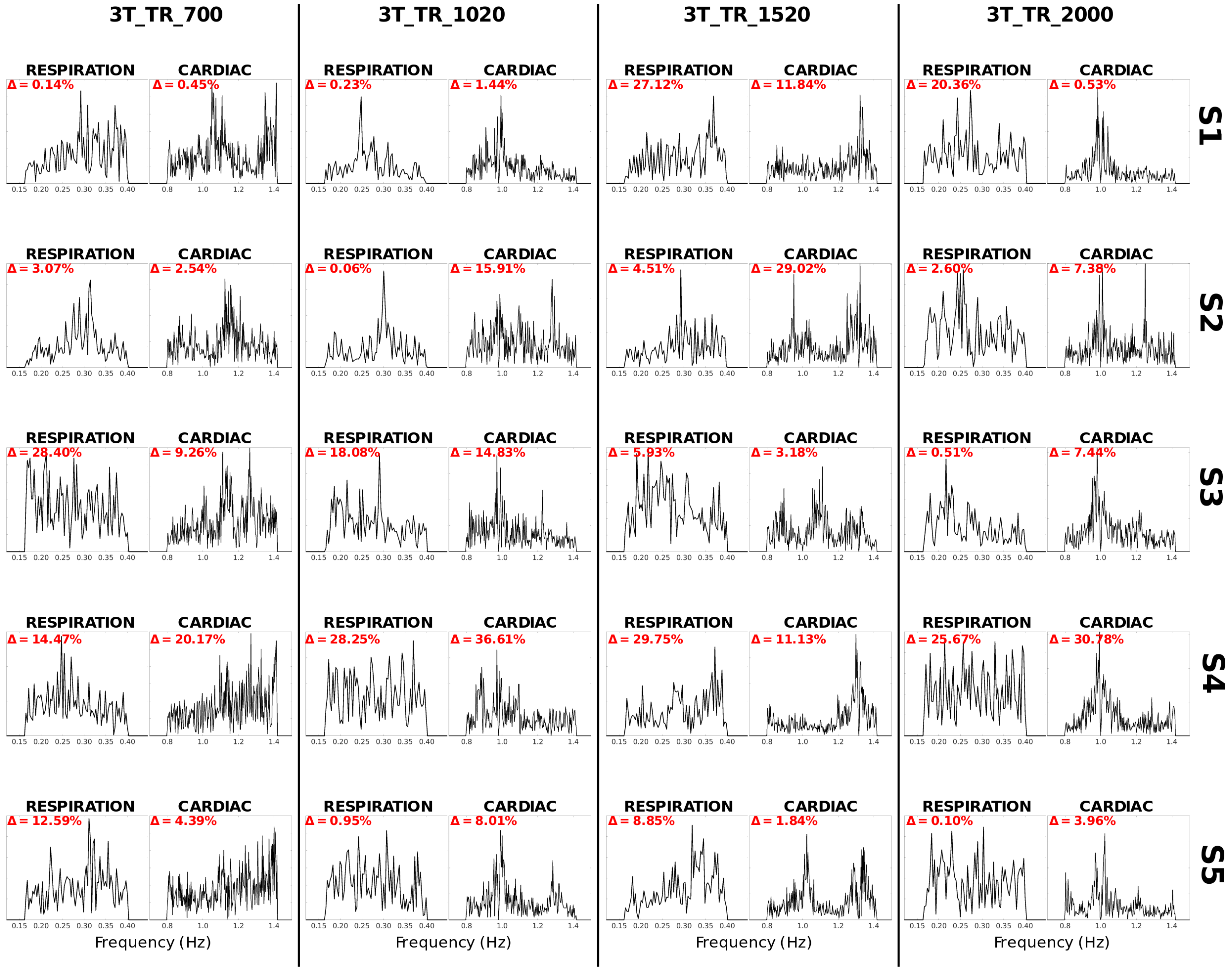


*sFig. 8: Power spectra of the PESTICA estimators (five first subjects of the 3 T study). Delta values indicate the deviation of the fundamental physiological frequencies to the expected values given by the external recordings.*

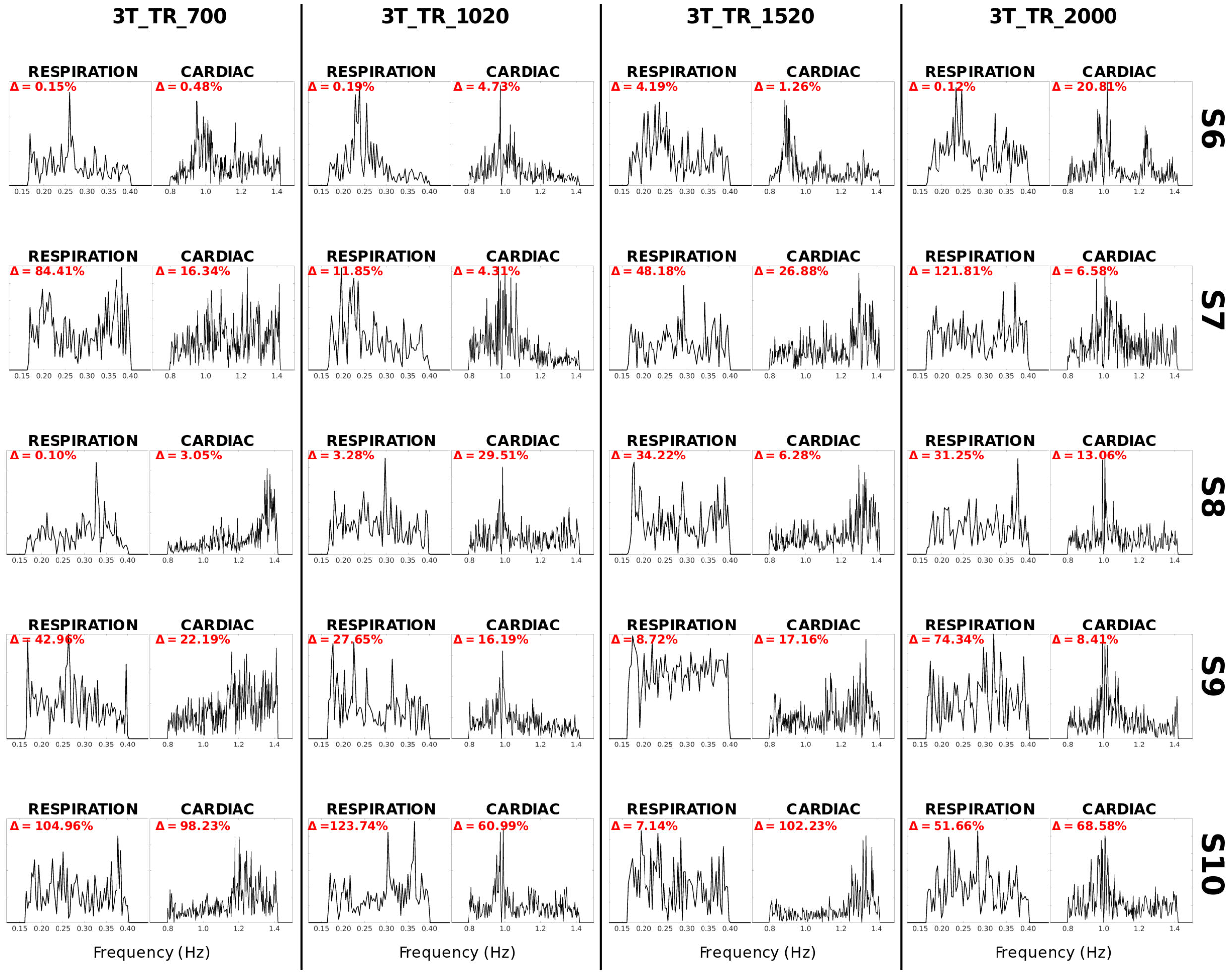


*sFig. 9: Power spectra of the PESTICA estimators (five last subjects of the 3 T study). Delta values indicate the deviation of the fundamental physiological frequencies to the expected values given by the external recordings.*
